# Enhancing Tumor Perfusion and Nanomedicine Delivery via Endogenous Nitric Oxide Release by Methyl Palmitate Nanoparticles

**DOI:** 10.64898/2026.03.02.709151

**Authors:** Roberto Palomba, Elizabeth Isaac, Raffaele Spanò, Federica Piccardi, Benedict McLarney, Elana Apfelbaum, Nermin Mostafa, Charlene Hsu, Jan Grimm, Paolo Decuzzi

**Affiliations:** Laboratory of Nanotechnology for Precision Medicine – Fondazione Istituto Italiano di Tecnologia, Via Morego 30, 16163, Genova, Italy; Molecular Pharmacology Program, Memorial Sloan Kettering Cancer Center, 1275 York Avenue, New York, NY, USA; Department of Radiology, Memorial Sloan Kettering Cancer Center, 1275 York Avenue, New York, NY, USA; School of Medicine/Division of Oncology, Center for Clinical Sciences Research, Stanford University, 269 Campus Drive, Stanford, CA 94305 - USA

**Keywords:** tumor perfusion, vascular permeability, nitric oxide, methyl palmitate, nanomedicine

## Abstract

Despite a few clinical successes, the efficacy of cancer nanomedicines remains limited by rapid clearance by the mononuclear phagocytic system and poor permeation across the abnormal tumor vasculature. We previously showed that methyl palmitate nanoparticles (MPN) can safely and reversibly inhibit the phagocytic activity of immune cells for several hours, thereby improving tumor accumulation and the efficacy of systemically administered nanomedicines. Here, we demonstrate that, on a shorter time scale, MPN can induce vasodilation, introducing an additional mechanism to enhance the accumulation of therapeutic agents within the malignant tissue. Upon internalization by macrophages and endothelial cells, MPN trigger the release of endogenous nitric oxide (NO), a key mediator of vasodilation, in a concentration-, and time-dependent manner. Following MPN administration, raster-scanning optoacoustic mesoscopy (RSOM) revealed vasodilation across multiple tissues, with the strongest effect observed in tumors. To assess enhanced tumor accumulation, we injected 70 kDa fluorescent dextran and demonstrated via histology a markedly increased fluorescence signal exclusively in MPN-treated tumors compared to controls 24 hours later. In addition, positron emission tomography (PET) imaging of ^89^Zr-labeled clinical iron oxide nanoparticles (Feraheme) showed significantly greater tumor accumulation after a 15-minute MPN pretreatment. Finally, general serum biochemistry panels and histological analyses of major organs in healthy mice revealed no toxicity following either single or repeated MPN dosing. Overall, this study demonstrates that MPN-induced vasodilation occurring within minutes enhances intra-tumoral deposition of macromolecules and small nanoparticles. Together with their longer-term effects on phagocytosis inhibition, these findings indicate that MPN can improve therapeutic delivery through complementary, time-dependent mechanisms that increase tumor perfusion and vascular permeability.

## Introduction

The delivery of macromolecules and nanoparticles to tumors relies on a complex series of processes occurring at both the systemic and tissue levels. At the systemic level, Kupffer cells positioned as sentinels along the vascular lumen of liver sinusoids, splenic macrophages forming a dense filtration network, alveolar macrophages lining the pulmonary microvasculature, and circulating monocytes collectively contribute to the rapid removal of foreign materials, including therapeutic nanoparticles (e.g., nanomedicines)^1–4^. At the tissue level, mass transport from the vascular compartment into the tumor parenchyma is impaired by the dysregulated and abnormal tumor vasculature, which leads to poor blood perfusion and irregular endothelial fenestrations, thereby limiting the intra-tumoral accumulation of systemically injected therapeutic agents^5^. Blood flow to tumors is further restricted by the immature vasculature bed ^6^, which is rapidly outgrown by the tumor cells itself. As a result, some vessels collapse under elevated interstitial fluid pressure^7^, while others remain structurally undeveloped, contributing to increased vascular resistance^8^. Together, these factors result in reduced and highly heterogeneous perfusion, ultimately limiting the transport of small molecules, macromolecules as well as nanoparticles from the bloodstream into the tumor tissue and their accumulation within the bulk tumor.

In previous work, we demonstrated that systemically administered methyl palmitate nanoparticles (MPN) can transiently and safely suppress the phagocytic activity of macrophages, thereby prolonging the circulation time and enhancing the tumor accumulation of systemically injected nanomedicines^2^. Different strategies have been recently explored by other groups to improve intra-tumoral transport of systemically administered therapeutic agents by acting at tumor level. One such approach focuses on remodeling and normalizing the tumor vasculature using anti-angiogenic therapies^9,10^. By partially restoring vascular architecture, these treatments aim to improve tissue perfusion. However, vascular normalization often reduces endothelial fenestration size, which can limit the intra-tumoral accumulation of most conventional systemically injected nanomedicines^11^. More recently, immunomodulatory strategies have sought to exploit the tumor immune landscape by activating immunostimulatory cells to combat cancer. These approaches can also indirectly affect the tumor vasculature, through cytokine signaling within the tumor microenvironment^10^. Although such strategies may enhance vascular function, their effects typically require days to develop, as they depend on extensive cellular and biophysical remodeling of the malignant tissue. More recent efforts have shifted toward strategies that exploit the existing, although abnormal, tumor vasculature to achieve more immediate improvements in blood perfusion. One such strategy employs focused high-intensity ultrasound in combination with microbubbles to transiently increase vascular permeability, followed by the administration of therapeutic agents^12^. However, to date, this method has primarily resulted in perivascular nanoparticle accumulation without a clear increase in tumor perfusion. Another promising strategy centers on modulating nitric oxide (NO) signaling. NO is a potent mediator of rapid vasodilation^13^ and leveraging this effect could improve the short-term transport of macromolecules and nanoparticles by widening the blood vessels^13^. Despite this promise, current NO-based therapies face significant practical challenges. The high chemical reactivity and difficult handling of gaseous NO have led researchers to develop NO donors; however, most existing formulations suffer from limited biocompatibility, suboptimal biodegradation profiles, or unfavorable release kinetics^14^.

To overcome these limitations, we developed an alternative strategy that stimulates the release of endogenous NO directly within vascular endothelial cells and macrophages, rather than delivering NO via exogenous agents. This approach is enabled by the methyl palmitate nanoparticles (MPN), which are fabricated from naturally occurring components: albumin, the most abundant serum protein, serves as the structural scaffold; and methyl palmitate, an endogenous lipid, functions as the bioactive payload. In the present study, we investigate an additional and previously unexplored property of MPN: their ability to modulate vascular tone and influence the tumor vascular bed, thereby improving the delivery of systemically injected molecules and nanomedicines. This work is motivated by recent findings identifying methyl palmitate as a potential mediator of vasodilation in retinal tissue^15^, possibly through NO-dependent mechanisms^16,17^. Given the absence of prior reports examining vasodilation with nanoparticle-formulated methyl palmitate, we completed initial *in vitro* studies to measure the capacity of MPN to induce NO production in macrophages and endothelial cells. We then assessed the vasodilatory activity of MPN *in vivo* by measuring real-time changes in tumor vessel tone following systemic MPN administration. Finally, we examined whether these vascular effects translated into improved transport and intra-tumoral deposition of macromolecules and nanoparticles.

## RESULTS AND DISCUSSION

### Physico-chemical characterizations of methyl palmitate nanoparticles and cellular uptake

Methyl palmitate nanoparticles (MPN) were produced by nanoprecipitation, following a procedure described in our previous publication^2^. Briefly, an ethanol-based methyl palmitate solution was mixed and sonicated with an albumin solution in PBS. After purification, MPN exhibited a hydrodynamic diameter of ∼200 nm (d = 185.06 ± 0.68 nm), a narrow size distribution (PDI = 0.097 ± 0.019), and a negative surface electrostatic potential (ζ = -31.5 ± 0.306), as summarized in **Figure 1a**. MPN appeared predominantly spheroidal in shape, as documented by the transmission electron microscopy image in **Figure 1b**, and its inset. Given their morphology and based on observations from previous studies,^18–20^ MPN are expected to be efficiently internalized by resident macrophages. To this end, a fluorescent version of MPN was generated by dispersing lipid-Cy5 molecules with methyl-palmitate in the original ethanol solution. The resulting Cy5-MPN displayed physicochemical features comparable to unlabeled MPN; a detailed physicochemical characterization of Cy5-MPN is available in **SI1**. To test MPN internalization, bone marrow-derived macrophages (BMDM), RAW264.7 cells, and human umbilical vein endothelial cells (HUVEC) were cultured and incubated with Cy5-MPN for 16 hours. Cells were then fixed, stained and observed under a confocal microscope **(Figure 1c**). Fluorescent MPN were predominantly observed in a perinuclear position across all cell types, suggesting efficient cellular uptake. The intracellular localization of MPN was further confirmed via transmission electron microscopy of BMDM (**Figure 1d**), following a 4-hour incubation with the nanoparticles. Samples were counterstained to visualize both lipid and protein components, which constitute the MPN structure. Discrete, spheroidal electron-dense structures surrounded by a darker boundary were observed exclusively in MPN-treated cells. This darker rim is consistent with an albumin-rich outer layer, as albumin contains sialic acid residues whose carboxyl groups are highlighted by uranyl acetate staining. The preservation of particle shape and their appearance as individual entities, rather than aggregates, suggest that the nanoparticles were internalized shortly before fixation. No such structures were detected in untreated control cells.

**Figure 1.**
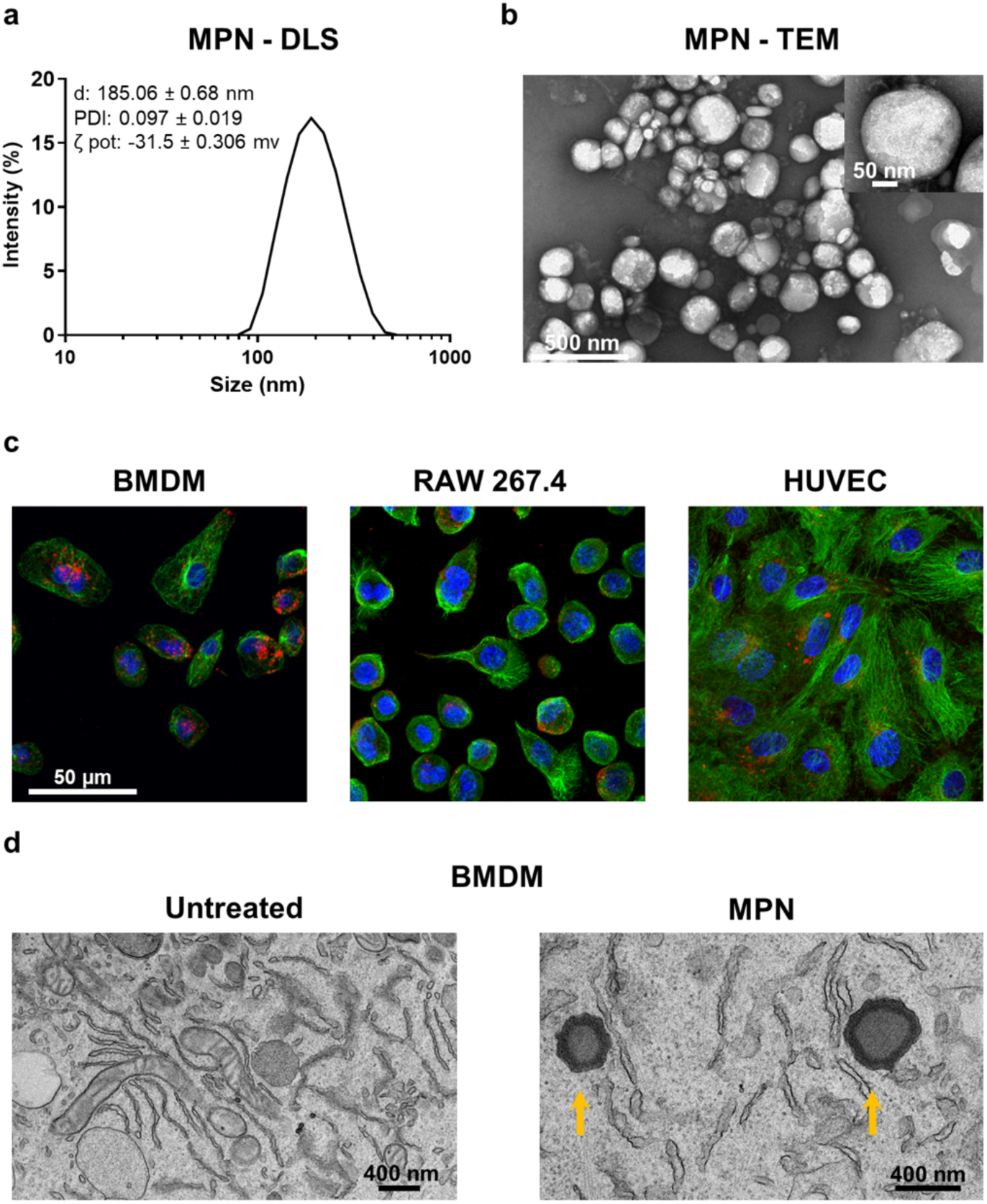
Physicochemical characterization and cellular uptake of MPN. **a.** Dynamic light scattering (DLS) analysis showing the hydrodynamic size distribution of MPN, including diameter (d), polydispersity index (PDI), and surface electrostatic potential (ζ). **b.** Transmission electron microscopy (TEM) images of MPN showing their spheroidal morphology (inset: higher magnification). **c.** Confocal microscopy images demonstrating MPN internalization in BMDM, RAW 264.7 cells, and HUVEC (red: MPN; blue: nuclei; green: tubulin in BMDM and actin in RAW 264.7 cells and HUVEC). **d.** TEM images of BMDM showing internalized MPN (yellow arrows).

### Nitric oxide release by immune and vascular cells exposed to methyl-palmitate nanoparticles

Methyl palmitate was recently identified as a highly potent endogenous vasodilator released by perivascular adipose tissue and retinal tissues, acting on the vessels of the aorta and retina, respectively^15,16^. Despite these findings, the mechanisms underlying its vasoactive function remain debated. Given the poor bioavailability of free methyl palmitate and the impracticality of systemic administration of the lipid alone, we investigated whether methyl palmitate nanoparticles (MPN), as carriers of methyl palmitate, could also exert vasodilatory effects. Upon systemic administration, MPN primarily encounter two major cell populations: endothelial cells lining the blood vessel walls and immune cells of the mononuclear phagocyte system (MPS)^21,22^. We hypothesized that exposure to and subsequent internalization of MPN could stimulate NO release in both immune and endothelial cells *in vitro*. To test this hypothesis, Griess assays were performed at predetermined time points on culture media collected from BMDM and HUVEC incubated with MPN. The release of NO was detected within minutes from MPN incubation, as shown in **Figure 2 a-d**, which presents the change in nitrite levels over time, for both cell types and varying the MPN concentration from 0 to 0.25 mM equivalent dose of methyl palmitate. Nitrite levels peaked around 10 min and remained relatively stable for up to 1 hour. Then, nitrite concentrations gradually declined for the BMDM, while they remained nearly constant for the HUVEC up to 8 hours. For both cell types, nitrite production increased in a concentration-dependent manner with increasing MPN dose. Because NO release and synthesis are calcium-dependent processes, we then investigated changes in intracellular calcium concentration in BMDM during the first few minutes after MPN incubation. Notably, a correlation between Ca^2+^ concentration and NO release has also been reported in the case of HUVEC by other researchers who linked it to the mobilization of Ca^2+^-ions from endoplasmic reticulum stores, influencing both NO release and production^23,24^. Accordingly, we used time-lapse fluorescence microscopy to monitor intracellular Ca²⁺ levels using the calcium-sensitive dye Fluo-4 AM. BMDM were seeded in microscopy chamber slides, loaded with Fluo-4 AM, and exposed to MPN immediately before image acquisition. During the first few minutes, the intracellular Ca^2+^ levels steadily increased to reach a maximum around 10 min (**Figure 2e**). Selected frames from the time-lapse recordings are shown in **Figure 2f** for three different representative cells. The fluorescent intensity measured in the region of interest (ROI) associated with the cells increases with time documenting the progressive rise in intracellular Ca^2+^ concentrations. Notably, individual cells exhibited different rates of Ca²⁺ increase, but all tended to reach comparable levels within the first 10 minutes.

**Figure 2.**
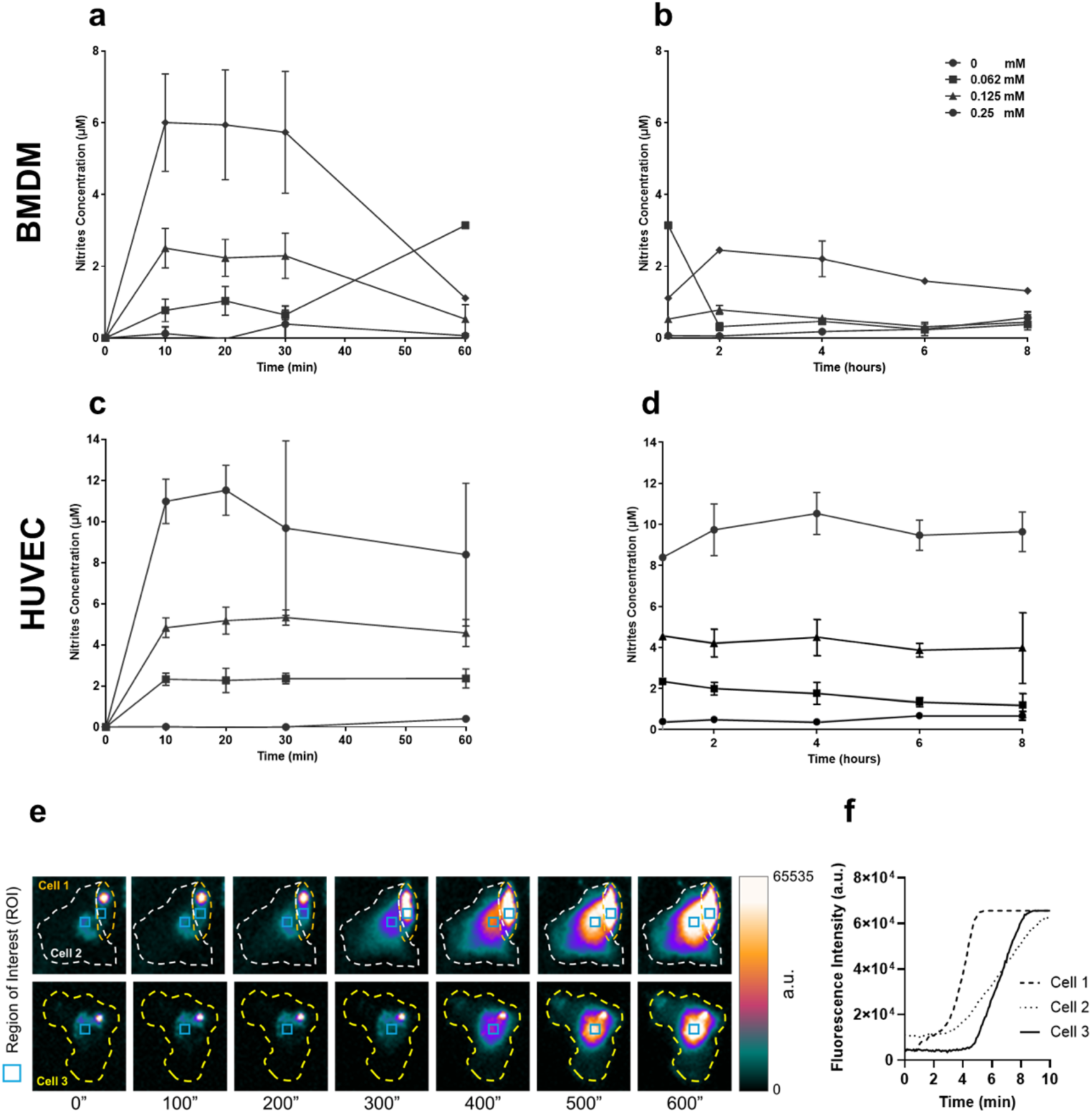
Nitric oxide release and calcium signaling in response to MPN treatment. **a-d,** Griess assay measuring nitrite accumulation in culture media of BMDM and HUVEC following incubation with increasing doses of MPN at different time points. **e,** Time course of intracellular Ca²⁺ concentration in BMDM measured by live-cell fluorescence imaging immediately after exposure to MPN. **f,** Representative single-cell Ca²⁺ responses observed by time-lapse microscopy following MPN exposure. (white and yellow dashed lines identify the entire cell body; azure boxes identify the region of interest (ROI) for Ca²⁺ level measurements as in 2b.

### Analysis of vasodilation in healthy and tumor tissues induced by methyl palmitate nanoparticles

We next investigated whether methyl palmitate retains its vasoactive function when administered *in vivo* as a nano-formulation. Initial qualitative observations in mice following the systemic administration of MPN revealed a rapid and pronounced vasodilatory response in the ear vasculature of mice, occurring within 5-15 minutes post-injection **(Figure 3a)**. MPN-injected mice displayed enhanced visibility of blood vessels and fine capillary networks, accompanied by increased ear redness. In contrast, mice injected with an equal volume of the vehicle (PBS), lacked as discernible capillary networks in the ears.

**Figure 3.**
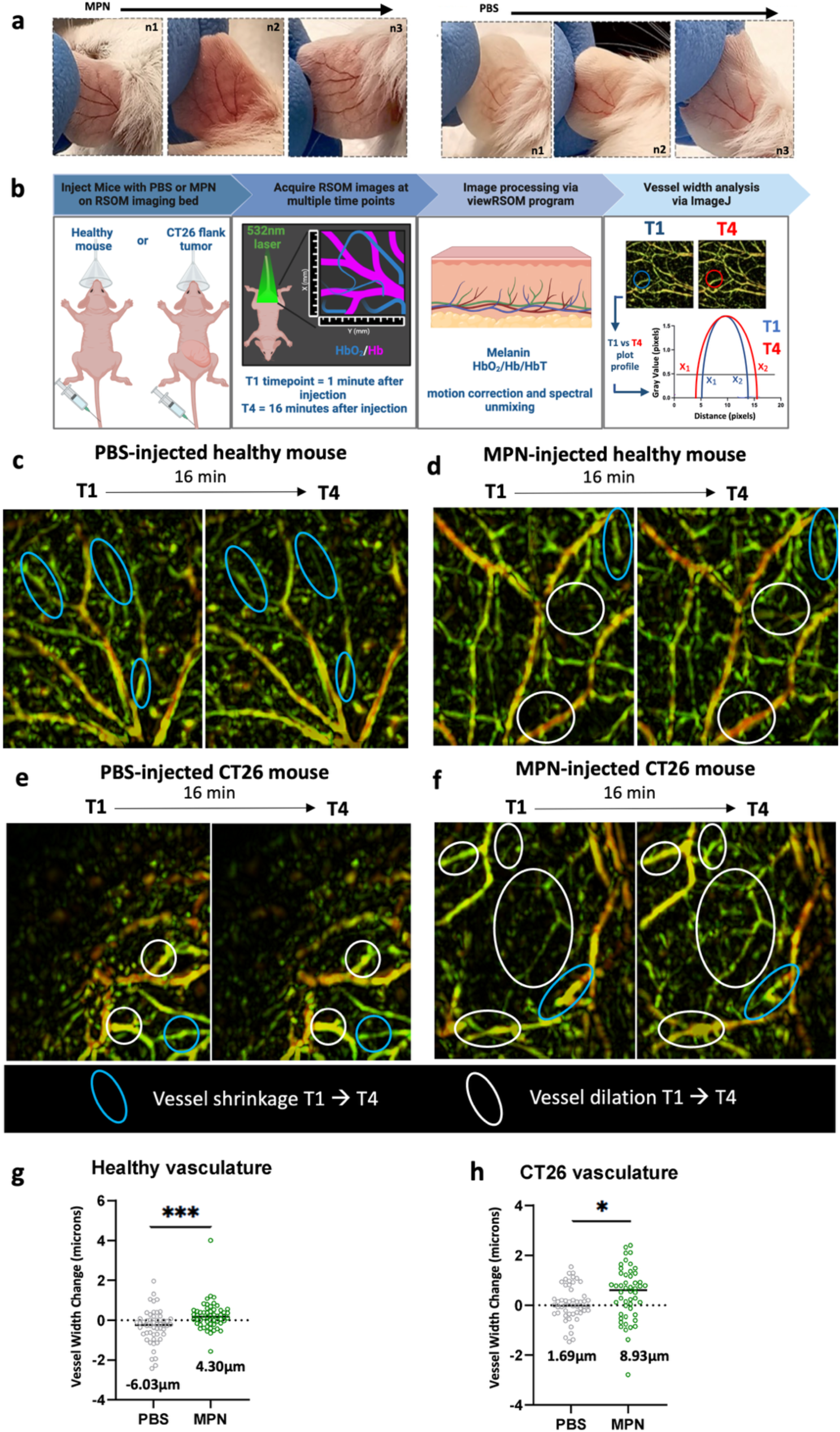
Systemic MPN injection induces varying levels of vasodilation in normal and tumor tissues vasculature. **a,** pronounced vasodilation and redness is observed in mouse ears within 5-15 minutes of systemic MPN injection, compared to systemic PBS-injection. **b,** RSOM schematic for vessel width analysis. Non-tumor-bearing mice and mice implanted with flank CT26 tumors were anesthetized on the RSOM imaging bed and injected with MPN or PBS. RSOM images of healthy tissue and tumors tissue vasculature were captured one-minute post-injection (T1) up till 16 minutes post-injection (T16). After motion correction and spectral unmixing of RSOM acquired images, the change in vessel width of approximately 17 vessel regions per experimental group (n=3) was analyzed using imageJ. Representative T1 vs T4 RSOM images of vasculature in **c,** PBS-injected healthy mice **d,** MPN-injected healthy mice **e,** PBS-injected CT26 tumor-bearing mice and **f,** MPN-injected CT26 tumor-bearing mice. (Vessel regions circled in blue represent examples of vessel width reduction that occurred from T1 to T4. Vessel regions circled in white represent examples of vessel width expansion or dilation that occurred from T1 to T4)**. g,** quantification of change in vessel width in approximately 17 vessel regions from T1 to T4 in vasculature of healthy mice injected with PBS or MPN (n=3 mice, approximately 50 vessel regions analyzed per group). Average change in vessel width (µm) from T1 vs T4 for approximately 50 vessel regions is indicated in the graph. f, quantification of change in vessel width in approximately 17 vessel regions from T1 to T4 in vasculature of CT26 tumor-bearing mice injected with PBS or MPN (n=3 mice, approximately 50 vessel regions analyzed per group). Average change in vessel width (µm) from T1 vs T4 for approximately 50 vessel regions is indicated in the graph, for all experimental groups. An unpaired t-test was used to assess statistical significance. *** = p<0.001 and * = p<0.05.

To quantitatively assess vasodilation, we utilized raster scanning optoacoustic mesoscopy (RSOM) to image and analyze the healthy and neoplastic vasculature in response to systemic MPN administration. RSOM is a non-invasive, high-resolution optoacoustic imaging technique that uses laser pulses to excite tissue chromophores, primarily hemoglobin, within blood vessels. The resulting thermoelastic expansion generates ultrasound signals that are detected and reconstructed into detailed vascular images^25^. For this study, healthy nude mice and nude mice bearing CT26 colon carcinoma flank tumors were intravenously injected with either PBS or MPN, and RSOM images were acquired at short intervals (five minutes each) post injection. Four different experimental groups were evaluated: PBS-injected healthy (non-tumor-bearing) mice, PBS-injected CT26 tumor-bearing mice, MPN-injected healthy (non-tumor-bearing) mice, and MPN-injected CT26 tumor-bearing mice. For all experiments, a dose corresponding to 1.875 mg of methyl palmitate per mouse was administered intravenously. Because MPN are resuspended in PBS, an equivalent volume of PBS was used as a vehicle control injection (negative control). Of note, as previously mentioned, methyl palmitate alone cannot be injected intravenously due to its poor solubility. Nude mice were used for this study to avoid depilation creams in the imaging area, which can cause epidermal irritation, redness, and scarring that interfere with RSOM signal acquisition and vascular visualization.

RSOM images were acquired once before injection, 1-minute post injection (T1), and every 5 minutes up to 20 minutes post-injection (T2 = 6 min, T3 = 11 min, T4 = 16 min, and T5 = 20 min). Analysis of the post-acquisition images revealed that maximal vasodilation occurred approximately 16 minutes after MPN injection (T4), and this time point was therefore selected for downstream vessel width analysis. **Figure 3b** summarizes the steps of the RSOM experiment for quantifying vessel width changes in tumor and healthy tissues after mice were injected with PBS or MPN. Vessels were color-coded according to the frequency of the detected ultrasound signal, with larger and deeper vessels (33 – 99 MHz) appearing red, and smaller, more superficial vessels (11 – 33 MHz) appearing green ^26^. Final images were presented as maximum intensity projections (MIP) composed of merged red and green frequency sub-bands, resulting in composite green, yellow, and orange vascular structures that represent total hemoglobin content in the imaged region.

For vessel width quantification, 17 – 20 individual vessel regions were randomly selected from blinded images for each experimental group (n = 3 mice per group). The vessel widths at 16 minutes post injection (T4) were then directly compared to those of the same vessels at 1 minute post injection (T1). In total, approximately 50 vessel regions were analyzed for both healthy and CT26 tumor vasculature in mice injected with either PBS or MPN. ImageJ was used to generate T1 – T4 image stacks and perform vessel width measurements as described in the **Materials and Methods** section. In healthy mice, the vessel width increased to a greater extent between T1 and T4 in mice injected with MPN compared to those receiving PBS injection (**Figure 3c, d** and **g**). Although ∼50 vessel regions were selected over the entire scan volume from n=3 mice for quantitative analysis, **Figure 3c-f** displays RSOM images from one representative mouse per group with 3 – 5 vessels highlighted with blue or white circles, signifying vessel shrinkage or dilation, respectively. **SI2 and SI3** contain T1 and T16 RSOM images acquired from n=3 mice for each experimental group. In healthy mice that received the MPN injection, a 4.3 μm increase averaged across the approximately 50 vessel regions analyzed, occurred between T1 and T4 (**Figure 3g**). In contrast, in mice injected with PBS, the vessel analysis indicated a 6.03 μm *reduction* in width, occurring between T1 and T4 (**Figure 3g**). One explanation for this observation is that in PBS injected mice, an initial volumetric expansion in vessel width 1-minute post-injection (T1) occurs, which dissipates at 16 minutes (T4) post injection, resulting in the measurement of a negative vessel width from T1 to T4. It is possible that compensatory vasoconstriction could have occurred as well. In MPN-injected mice, there may also be an initial volumetric expansion post injection at the T1 timepoint, but since MPN possesses vasodilative effects, sustained vasodilation is likely observed at T4.

A similar but more pronounced effect was observed in CT26 tumor–bearing mice. Tumor vessels in MPN-injected animals displayed significantly greater vasodilation than those in PBS-injected controls (**Figure 3e, f** and **h**). The average vessel width changes between T1 and T4 was +8.93 μm on average in MPN-injected mice across the 50 vessel regions analyzed, while PBS-injected mice only displayed a +1.69 μm on average increase (**Figure 3h**). Interestingly, MPN injection in both the healthy and tumor vasculature resulted in the appearance of vessels at timepoint T4, that were too difficult to capture via RSOM imaging at timepoint T1. Some striking examples are identified with white circles in **Figures 3d** and **3f**. These observations indicate that, in addition to dilating larger pre-existing vessels, MPN induce vasodilation of smaller vessels, rendering them detectable by RSOM imaging. Importantly, one key difference between vascular responses of healthy and tumor tissues is that MPN-mediated vasodilation was more pronoucned in tumor tissues, resulting in a two-fold greater vasodilation effect in tumor vasculature compared to healthy skin tissue vasculature (8.93 micron increase in tumor tissue vasculature compared to a 4.30 micron increase in non-tumor tissue vasculature).

### MPN-induced vasodilation enhances the intra-tumoral deposition of macromolecules

After demonstrating that MPN induce tumor vessel dilation, we next investigated whether this vascular priming effect could lead to a measurable increase in the intra-tumoral accumulation of macromolecules when administered systemically after MPN injection. To this end, CT26 tumor bearing mice were first injected intravenously with MPN and, 15 – 20 minutes later, retro-orbitally injected with 70 kDa fluorescent dextran, as outlined in **Figure 4a**. Twenty-four hours after injection, mice were sacrificed, tumors were explanted, and tissue sections were prepared for fluorescence analysis. Fluorescence microscopy imaging was used to compare dextran accumulation (green signal) within tumors from mice which were pretreated (primed) with PBS or MPN. Quantitative analysis was performed by measuring the total tumor slice area and the fraction of that area positive for dextran fluorescence. In **Figure 4b**, two representative CT26 tumor sections were reported, one for the PBS injection group and one for the MPN pretreated group. For each section, the full area used for quantification is shown on the left, and the dextran-positive area is shown on the right. By comparing the positive areas of the slices of the two groups (PBS and MPN), a greater extent of fluorescent dextran accumulation is clearly observed in tumors from mice pretreated with MPN. These two representative images were false-colored to further highlight the difference in dextran fluorescence intensity and topographic distribution within the CT26 tumors (**Figure 4c)**. In PBS injected mice, 70 kDa dextran was largely confined to peripheral tumor regions and failed to penetrate the tumor core. In contrast, in the MPN pretreated mice, a more extensive diffusion of the fluorescence signal throughout the entire tumor slice is observed, indicating that the 70 kDa fluorescent dextran is able to deeply diffuse inside the core of the tumor while reaching higher intensities at its periphery. To compare dextran accumulation between PBS and MPN priming conditions, the dextran-positive area fraction of each slice was quantified. A statistically significant difference was observed between the two groups, with slices from MPN pretreated mice showing approximately 1.5-fold larger dextran-positive areas. These results indicate that short-term MPN priming is effective in enhancing macromolecule accumulation within solid tumors. Taken together, the data presented in **Figures 3** and **4** indicate that the transient and reversible vasodilation induced by MPN occurs preferentially at tumor lesions and promotes enhanced intra-tumoral accumulation and retention of macromolecules (such as 70 kDa dextran) for at least 24 hours post-injection.

**Figure 4.**
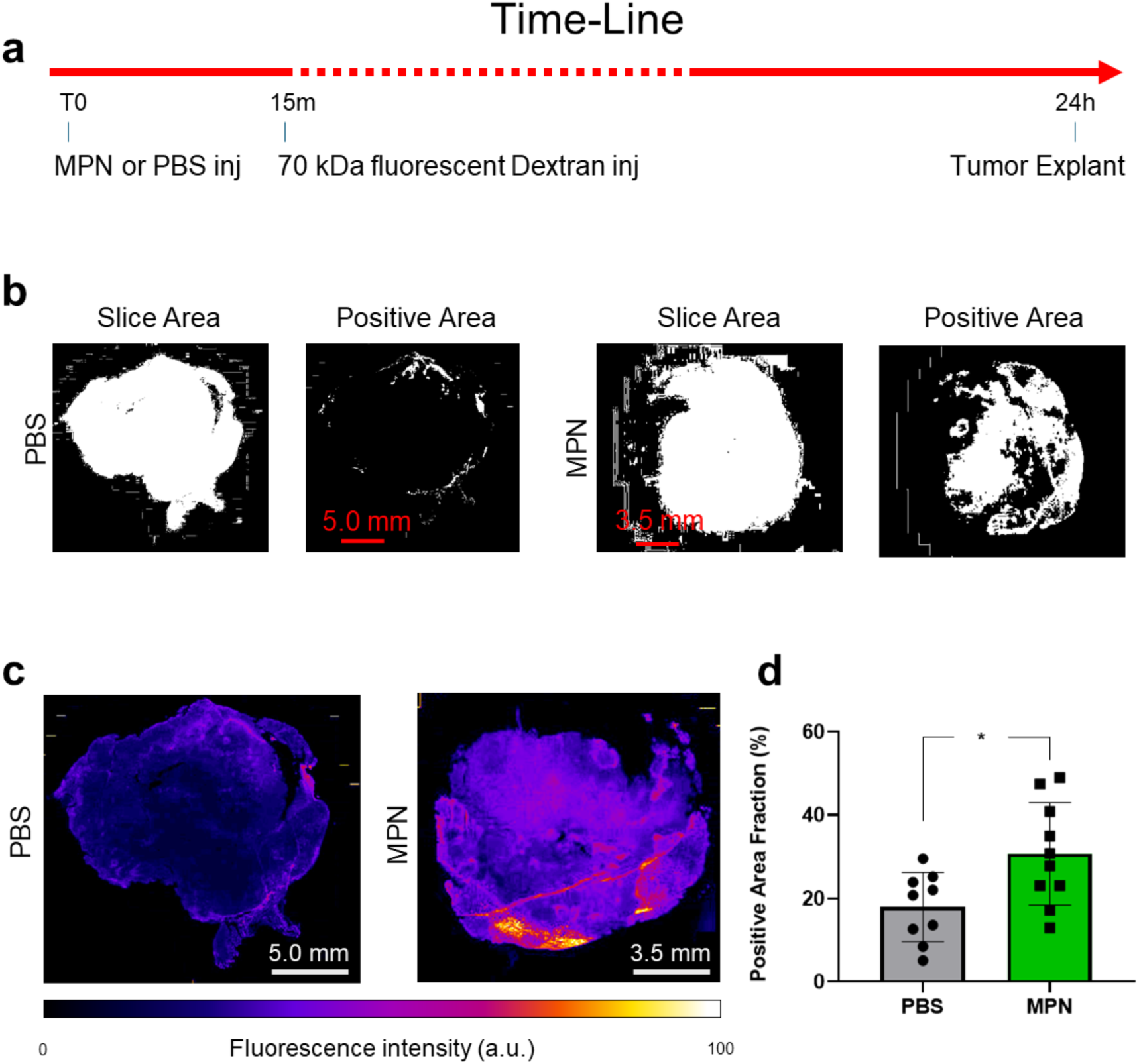
Quantification of intratumoral dextran accumulation. **a,** Timeline of treatments and procedures in CT26 tumor-bearing mice. **b,** Representative tumor slice area (left) and dextran-positive area (right) in PBS-injected and MPN-pretreated mice. **c,** False-colored representative tumor images showing spatial distribution and intensity of dextran fluorescence in PBS-DEX and MPN-DEX mice. **d,** Quantification of dextran-positive area fraction in tumor slices from PBS-DEX and MPN-DEX groups (p = 0.018).

### MPN-induced vasodilation enhances the intratumoral deposition of nanoparticles

We next tested whether MPN vascular priming could also improve the accumulation of the FDA-approved iron oxide nanodrug Feraheme (FH). FH is a 750 kDa ultrasmall superparamagnetic iron oxide nanoparticle, with a colloidal diameter ranging from 17 to 31 nm and is used clinically for the treatment of iron-deficiency anemia. More recently, FH has attracted interest as an anticancer agent due to its ability to generate reactive oxygen species (ROS) and induce oxidative stress-mediated cancer cell death^27^. In addition, FH has been reported to induce a pro-inflammatory phenotype in macrophages, supporting its potential use as an immune-stimulating anti-tumor adjuvant ^28,29^. FH can also be non-covalently loaded with various therapeutic small molecules, such as doxorubicin, enabling effective nanoparticle-based delivery of small molecules to tumors^30^. Given its broad range of applications in cancer therapy, strategies that enhance FH accumulation within tumor masses could further improve its therapeutic potential as anti-cancer agent.

For this study, chelator-free radiolabeling of FH with the positron-emitting radionuclide ^89^Zirconium (^89^Zr-FH) was completed. Chelator-free radiolabeling of FH has previously been used to enable PET/CT-based in vivo visualization, tracking, and quantification of FH nanoparticle biodistribution in tumor and non-tumor tissues, without altering the physiochemical properties of the nanoparticles.^31,32^ Following existing protocols^31^, pH neutralized ^89^Zr was incubated with FH nanoparticles at 95°C for 2 hours, after which deferoxamine (DFO) was used to chelate unbound radionuclide, followed by Amicon filtration of ^89^Zr-FH. This process yielded a radiolabeling efficiency exceeding 90%, with instant thin-layer chromatography indicating that 91.5% ± 9% (n = 3) of the radioactivity was bound to the nanoparticle (**Figure 5a**).

**Figure 5.**
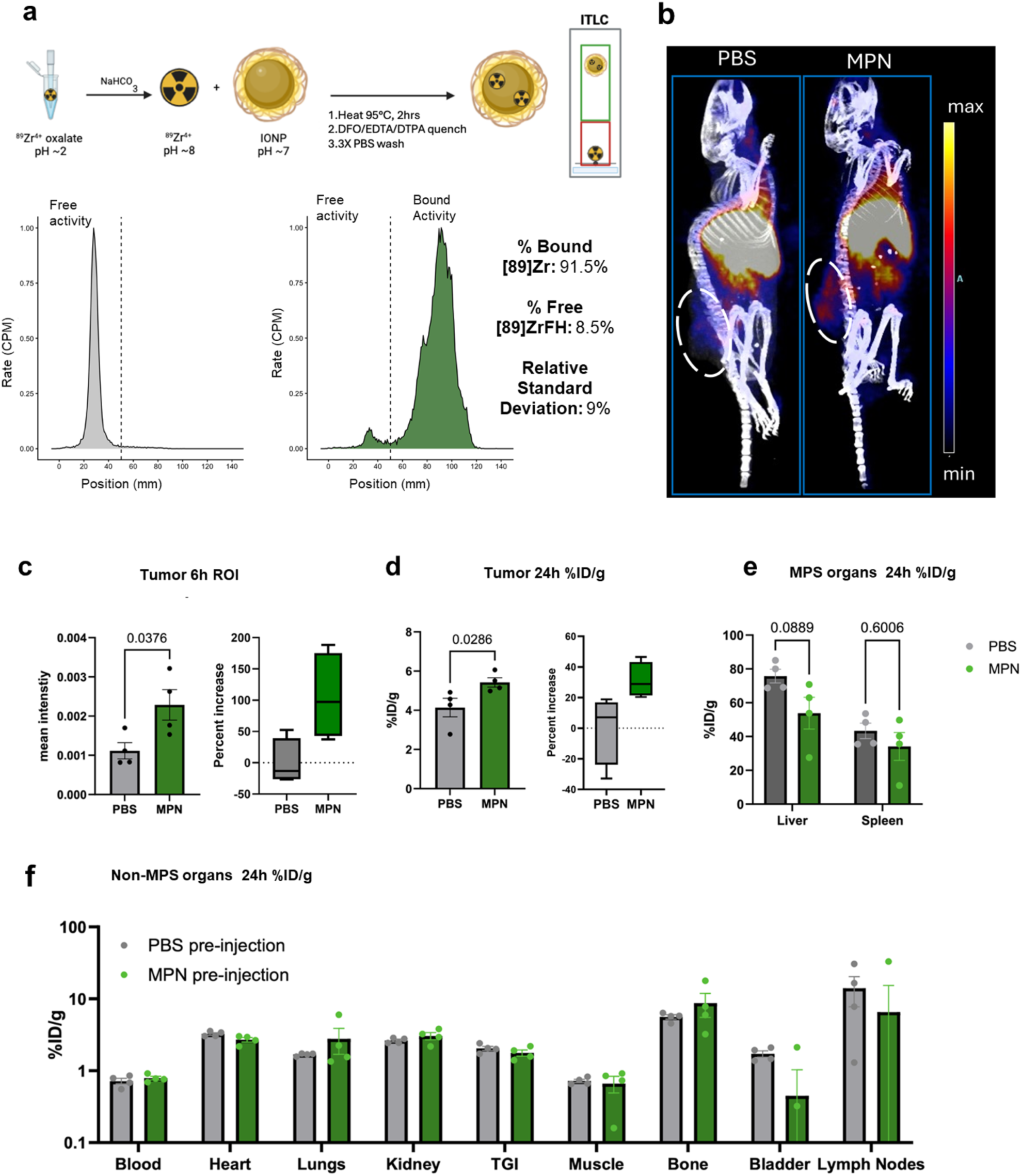
Quantification of intratumoral Feraheme (FH) accumulation. **a,** schematic of chelator free radiolabeling. FH is added to 89Zr-oxalate after neutralization with bicarbonate to a pH of ∼7-8. The solution is then heated to 95°C for 2 hours, after which unbound 89Zr is chelated with deferoxamine (DFO), followed by PBS washes in a 30kDa amicon filtration unit. To check radiolabeling efficiency, radiolabeled nanoparticle is run on an ITLC strip where free activity stays at the origin (corresponding to a large peak before the 50mm position on the ITLC strip) and radioactivity bound to the FH nanoparticle travels up the strip with the particle (corresponding to a large peak above 50mm and a small free activity peak below the 50mm position on the ITLC strip). **b,** PET/CT image displaying 89Zr-FH biodistribution 6-hours (h) after injection. Tissues with substantial nanoparticle uptake are labeled. **c,** 6h quantification of 89Zr-FH in tumor tissue (region of interest ROI) displayed as mean intensity and percent increase compared to PBS-preinjected mice. **d,** 24h analysis of % injected dose of 89Zr-FH per gram of tissue (%ID/g) in tumors. Percent increase compared to PBS-preinjected mice also displayed. **e,** Quantification of 89Zr-FH in macrophage-high organs of the mononuclear phagocyte system (MPS), displayed as %ID/g, at 24h post injection. **f,** Quantification of 89Zr-FH displayed as %ID/g, at 24h post 89Zr-FH injection in tissues with lower macrophage content compared to the liver and spleen.

*In vivo* biodistribution studies were then completed to determine if MPN pre-injection (*vascular priming*) could improve the accumulation of ^89^Zr-FH in CT26 tumor masses, compared to PBS (control) pre-injection. Mice were injected with 100µL of ^89^Zr-FH, 15 to 20 minutes post MPN priming or PBS injection. PET/CT imaging was performed 6 hours post-injection using a Siemens Inveon scanner to evaluate whole-body biodistribution. Images were reconstructed as maximum intensity projections (MIP) using the Inveon Research Workplace software. As expected, ⁸⁹Zr-FH exhibited typical nanoparticle biodistribution, with prominent signal in the lungs and liver (**Figure 5b**). PET/CT images of all animals (n=4 mice per PBS or MPN pre-injection group) in this study are in **SI4**. Quantitative region-of-interest analysis of PET images revealed an approximately twofold (∼100%) increase in tumor-associated ⁸⁹Zr-FH signal in MPN-pretreated mice compared with PBS-pretreated controls at 6 hours post-injection (n = 4 mice per group; **Figure 5c**).

Mice were subsequently sacrificed 24 hours after ⁸⁹Zr-FH injection, and organs were harvested for gamma counting to determine percent injected dose per gram of tissue (%ID/g). Tumors from MPN-pretreated mice exhibited a sustained ∼30% increase in ⁸⁹Zr-FH accumulation compared with PBS controls at 24 hours (**Figure 5d**), indicating prolonged retention following enhanced early deposition. Accumulation of ⁸⁹Zr-FH in the liver and spleen was also evaluated, since these organs contain high levels of tissue resident macrophages. Although MPN pretreatment showed a modest trend toward reduced FH accumulation in these organs, no statistically significant differences were observed (**Figure 5e**). This result is consistent with the relatively minor vasodilatory effects of MPN in healthy tissues and with the short (15-minute) pretreatment interval used in this study, which is insufficient to induce macrophage phagocytic suppression in the liver (previously observed at ∼ 4 hours).²

Similarly, no significant differences in ⁸⁹Zr-FH accumulation were detected in organs with lower macrophage density (**Figure 5f**). These findings align with RSOM analyses demonstrating that MPN-induced vasodilation is markedly greater in tumor vasculature than in healthy tissues. This tumor-selective vascular response likely reduces interstitial fluid pressure and enhances tumor perfusion, thereby facilitating increased nanodrug accumulation within malignant tissue.

Collectively, these results highlight the importance of pretreatment timing in exploiting the distinct biological effects of MPN. While longer pretreatment durations (∼ 4 hours) was previously reported to suppress macrophage uptake at the systemic level and favor delivery of larger nanoparticles (≥ 200 nm) ^2^, the shorter 15 minutes pretreatment used here preferentially enhances tumor perfusion, improving the accumulation of smaller nanoparticles (e.g., the 17 – 30 nm FH nanoparticles) and macromolecules (e.g., 70 kDa dextran).

### Toxicity tests on healthy methyl palmitate nanoparticles treated mice

To comprehensively evaluate the tolerability of MPN, we performed a series of toxicity assessments in healthy mice. First, we analyzed serological blood parameters in C57BL/6 mice that received a single high dose of MPN corresponding to 3.75 mg of methyl palmitate per 20 g body weight, administered via tail-vein injection. Blood samples were collected 24 hours after treatment. All measured serum chemistry parameters are summarized in **Table 1** and graphically represented in **SI5**. No statistically significant alterations were detected in any parameter except for uric acid, which was significantly reduced in MPN-treated mice compared with controls (**Table 1**). This reduction is consistent with increased nitric oxide (NO) production, as uric acid is a known endogenous scavenger of NO; unlike elevated uric acid levels, this decrease is not considered indicative of toxicity^33–35^. Hematological parameters measured under the same conditions likewise showed no significant differences between control and MPN pretreated mice (**Table 2** and **SI6)**.

To assess tolerability under repeated dosing, a separate cohort of mice received four injections of MPN administered twice per week over a 2-week period at a lower dose equivalent to 0.94 mg of methyl palmitate per 20 g body weight. Blood samples were collected 24 hours after the final injection, and serum chemistry results are listed in **Table 3** and **SI7**. Overall, no major abnormalities were observed. A mild increase in transaminases and serum cholinesterase was detected in MPN-treated mice; however, these changes did not reach statistical significance. A small but statistically significant increase in lactate dehydrogenase (LDH) was observed, although this elevation is not typically associated with clinical toxicity. All other parameters remained within physiological reference ranges. In contrast to the acute high-dose condition, uric acid levels in the repeated low-dose cohort showed a slight, non-significant increase, suggesting that higher MPN doses may be required to induce sufficient NO production to measurably reduce uric acid at the analyzed time point. Consistent with these findings, hematological analysis revealed no significant differences between MPN pretreated and control mice under repeated dosing conditions (**Table 4** and **SI8**).

To further evaluate potential tissue-level toxicity, additional mice were treated under the same dosing regimens used for serum and hematological analyses. Twenty-four hours after a single high-dose MPN injection, or 24 hours after the final injection in the repeated low-dose cohort, mice were sacrificed and major organs were harvested for histological examination. Hematoxylin and eosin (H&E) staining of liver, spleen, lung, kidney, and heart sections revealed no observable pathological abnormalities in mice treated with either a single high dose (3.75 mg methyl palmitate per 20 g body weight) or repeated low-dose injections (0.94 mg per 20 g body weight), compared with untreated controls (**Figure 6a,b**). Collectively, these results indicate that MPN are well tolerated in healthy mice following both acute and short-term repeated administration.

**Figure 6.**
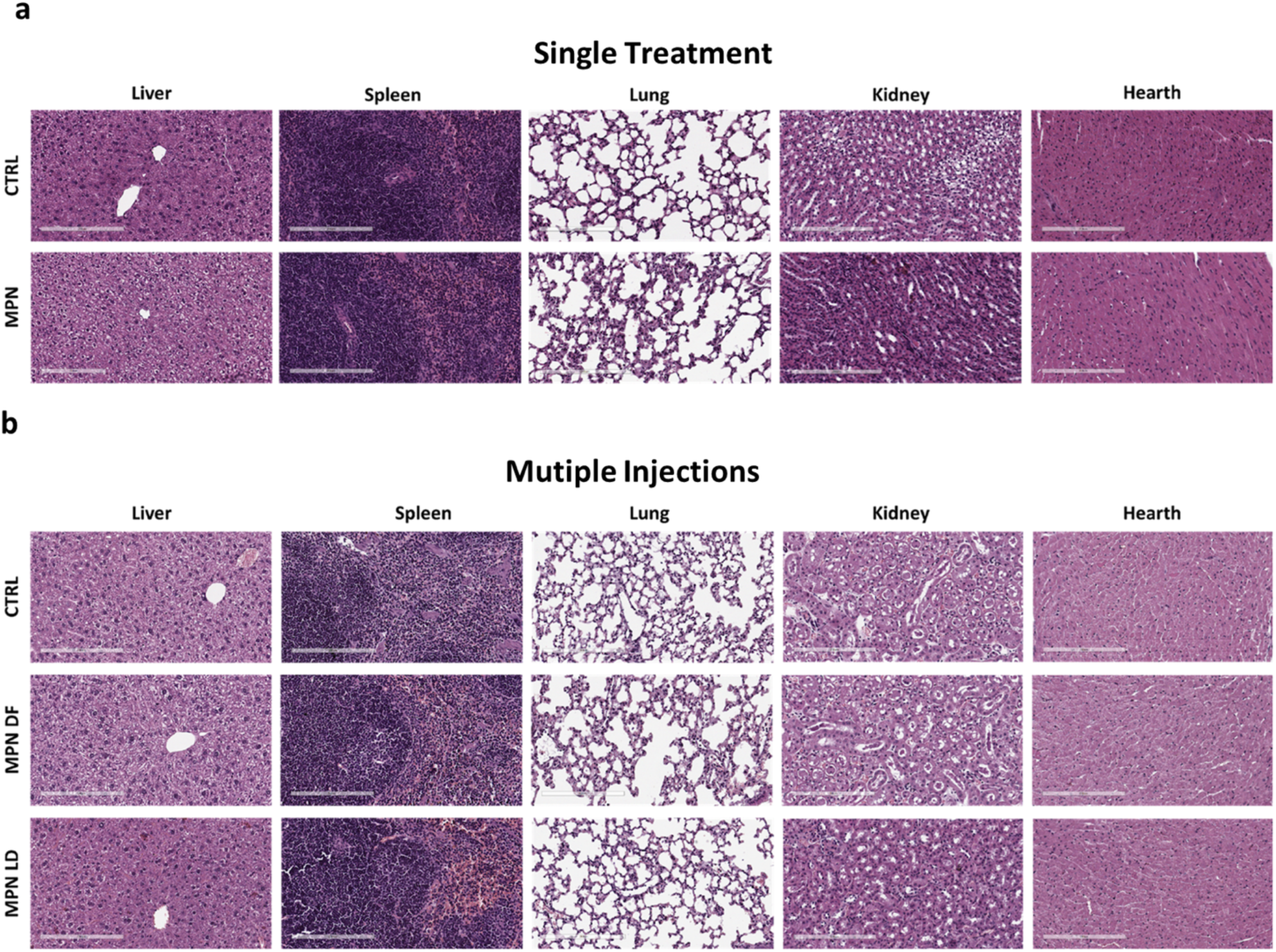
Toxicological evaluation of MPN in healthy mice. **a,** Serum uric acid levels measured 24 hours after a single intravenous injection of MPN (dose equivalent to 3.75 mg methyl palmitate per 20 g body weight). The reference range between the two dashed lines indicates values observed in 95% of the healthy population. **b,** Representative H&E-stained sections of major organs (liver, spleen, lung, kidney, and heart) from control mice and mice treated with a single high dose of MPN (3.75 mg methyl palmitate per 20 g body weight). **c,** Representative H&E-stained sections of major organs from mice treated with repeated MPN injections (twice per week for two weeks) at either full dose (3.75 mg per 20 g) or low dose (0.94 mg per 20 g). No overt histopathological abnormalities were observed.

## Conclusions

Our results demonstrate that MPN administration induces a rapid systemic vasodilation that is significantly more pronounced in tumor vasculature than in healthy tissues, including the highly vascularized skin. This early vasodilatory response, likely mediated by NO release from macrophages and endothelial cells, enhances vascular permeability and supports increased intratumoral uptake of macromolecules, such as 70 kDa dextran, and nanodrugs, such as the ∼ 20 nm Feraheme nanoparticles. Toxicology analyses further indicate that both single and repeated MPN dosing regimens are well tolerated in healthy mice.

Taken together with our previous findings, these data further suggest that the timing of MPN pretreatment can be strategically leveraged to optimize the delivery of therapeutic agents to tumors. Nanoparticles administered approximately 4 hours after MPN injection primarily benefit from reduced liver uptake, whereas those administered within ∼15 minutes of pretreatment benefit from enhanced tumor perfusion and extravasation. Because larger nanoparticles (> 100 nm) are more susceptible to clearance and less capable of passive extravasation, they may be better suited to the longer pretreatment window, while smaller nanoparticles (15 – 30 nm) and macromolecules (∼70 kDa) are ideally positioned to exploit the early vasodilation phase.

Finally, these complementary mechanisms may be combined. Sequential MPN priming or staggered delivery of therapeutics could enable simultaneous exploitation of both perfusion-driven enhancement and macrophage suppression, offering a flexible strategy to improve the delivery and efficacy of macromolecules and nanomedicines across a range of sizes and therapeutic modalities.

## Acknowledgments

Authors would like to acknowledge the Electron Microscopy Facility at IIT, particularly Roberto Marotta, Tiziano Catelani, Federico Catalano and Doriana Debellis. Authors would also like to thank the Nikon Imaging Center at IIT with a particular mention to Mattia Pesce, and the Animal Facility of “Fondazione Istituto Italiano di Tecnologia”, particularly Sara Morando and Monica Morini. Authors are also grateful to Andrea Contestabile, PhD for the support on the experiments aimed at measuring intracellular Ca^2+^ concentration and to Michele Schlich for the important feedback on the topic. Authors would also like to acknowledge the Molecular Cytology core at MSKCC for their assistance with tissue processing and image acquisitions, as well as the Animal Imaging Core for access to their PET/CT imaging equipment and image analysis software. Finally, we would like to acknowledge the partial support by the NIH (P30 cancer core support grant to S. Vickers, P30CA008748, R01 EB0033000; R01CA212379 and R01CA218615 to J. Grimm).

## Authors’ Contribution

**R.P.**: Design and preparation of MPN; internalization experiments by confocal microscopy; preparation of the samples for TEM; time-lapse microscopy experiment on calcium concentration; *in vivo* experiments about MPN tolerability; main organ histology; conceptualization, data curation, formal analysis, investigation, methodology, project administration, validation, visualization, writing original draft; review & editing. **E.I:** Completed mouse injections for RSOM experiments; completed RSOM image processing and data analysis; completed IV injection of MPN and retro-orbital injection of fluorescent dextran; harvested and fixed tissue for mounting and imaging; prepared radiolableded Feraheme and assisted with mouse injections; completed PET/CT imaging of mice injected with radiolabled Feraheme; completed PET/CT image processing and analysis; writing, reviewing, and editing of the manuscript **R.S.**: *in vivo* experiments about MPN tolerability, main organ histology. **F.P.**: *in vivo* experiments about MPN tolerability, main organ histology. **E.A.**: assisted with MPN injection for RSOM and RSOM image analysis. **B.M.**: gave orientation on how to use RSOM machine and plot profile tool of ImageJ for vessel width analysis. **N.M.**: assisted with mouse tail vein and retroorbital injections. **C.H.**: assisted with mouse tail vein injections and acquisition of RSOM images. **P.D.** and **J.G.**: project design and supervision, funding acquisition, review & editing.

## Materials and Methods

### Fabrication and characterization of Methyl Palmitate Nanoparticles

Methyl palmitate nanoparticles (MPN) were produced by using a self-assembly method: a solution 1:1 (V:V) of methyl palmitate (Merck – Sigma Aldrich – Germany) and ethanol containing 2 mg of methyl palmitate was added to 100 μl of a 50 mg/mL BSA (Merk - Sigma Aldrich - Germany) solution. The obtained mixture was sonicated for 1min into a water bath, washed in 1 mL of PBS (Thermo Fisher Scientific, USA) and centrifuged at 15,000 rpm at 4°C. Particles were washed and finally re-suspended in 1 mL of PBS or water (depending on the specific need) and sonicated for 1 minute. Average size, size distribution, and zeta potential of MPN were analyzed using dynamic light scattering. Samples were diluted with isosmotic double distilled water (1:100 v/v) to avoid multi-scattering phenomena and analyzed at 25 °C with a Zetasizer Nano (Malvern, U.K.), equipped with a 4.5 mW laser diode and operating at 670 nm as a light source, and the scattered photons were detected at 173 °C. A third order cumulative fitting autocorrelation function is applied to measure the average size and size distributions. The analysis was carried out according to the following instrumental setup: (a) a real refractive index of 1.59; (b) an imaginary refractive index of 0.0; (c) a medium refractive index of 1.330; (d) a medium viscosity of 1.0 mPa×s; and (e) a medium dielectric constant of 80.4. Transmission electron microscopy (TEM) micrographs were acquired using JEOL JEM 1011 (Jeol, Japan) electron microscope (Electron Microscopy Facility, Fondazione Istituto Italiano di Tecnologia, Genoa - Italy) operating with an acceleration voltage of 100 kV and recorded with a 11 Mp fiber optical charge-coupled device (CCD) camera (Gatan Orius SC-1000). MPN samples were diluted 1:100, dropped on 150-mesh glow discharged ‘Ultrathin’ carbon-coated Copper TEM grids, dried and directly observed.

### Cell Cultures

RAW264.7 cells were purchased from the American Type Culture Collection (ATCC, Rockville, MD, USA) and maintained in Dulbecco’s Modified Eagle’s Medium high-glucose (DMEM) (Euroclone - Italy) supplemented with 10% fetal bovine serum (ATCC) and 1% penicillin/streptomycin. Cells were grown at 37 °C in an 80% humid atmosphere of 5% CO_2_. Bone Marrow Derived Monocytes (BMDMs) from rats and from mice were isolated based on the following procedures. Briefly after specifying the animal, femurs were isolated, cleaned from surrounding tissues and washed in PBS (Thermo Fisher Scientific, USA), a cut was performed at both ends. PBS was used to flush the cavities, cells were harvested and plated in media supplemented with macrophage Colony-Stimulating Factor (mCSF) (10 ng mL−1) (Merk - Sigma Aldrich - Germany). The procedures were conducted following the guidelines of the Institutional Animal Care and Use Committee of IIT and the National Institutes of Health guide for the care and use of Laboratory animals (NIH Publications No. 8023, revised 1978). Culturing medium was changed after 3 days to remove unattached cells. BMDMs were used on the following day. Culture media: DMEM supplemented with 15% FBS, 1% penicillin/streptomycin, and rat M-CSF (Merk - Sigma Aldrich - Germany) (according to vendor indications).

Human Umbilical Vein Endothelial Cells (HUVEC) were cultured using Vascular Cell Basal Medium supplemented with 0.2% bovine brain extract, rh EGF (5ng/mL), L-glutamine (10mM), heparin sulfate (0.75 Units/mL), hydrocortisone (1µg/mL), ascorbic acid (50µg/mL), 2% fetal bovine serum, and 1% penicillin/streptomycin. CT26 Cells were cultured in RPMI base media supplemented with 10% fetal bovine serum and 1% penicillin/streptomycin. Cells were cultured under controlled environmental conditions (37 °C in 5% CO2).

### Particle internalization analysis via Confocal Microscopy

For cells imaging 65,000 BMDM, 65,000 RAW 267.4, and 6,500 HUVEC, were separately seeded into each well of a µ-Slide 8 well high (ibidi GmbH – Germany) maintaining culturing conditions, as described above. Cells were treated overnight with Cy5-MPN. After these treatments media was removed and cells were washed in PBS (Thermo Fisher Scientific, USA). Fixation was performed using a 3.7% solution of paraformaldehyde (Sigma Aldrich, USA) for 10 minutes; 3 washes with PBS (Thermo Fisher Scientific, USA) were performed after cell fixation. To highlight cell body BMDM were immunostained for alpha tubulin, while RAW 267.4 and HUVEC were stained for actin. for BMDM α-Tubulin antibody (Sigma - T5168) was used as primary antibody and Abcam - ab6786 was used as secondary antibody according to vendors indications. Actin was stained using Alexa Fluor 488 Phalloidin (Thermo Fisher Scientific, USA) according to vendor indication. For all the analyses, nuclei were stained using DAPI (Thermo Fisher Scientific, USA) following vendor indications. A 63× objective was used and a z-stack series was acquired (≥ 12 steps of 1,000 nm each were acquired per image).

### Transmission Electron Microscopy Analysis of BMDM

Transmission electron microscopy (TEM) micrographs were acquired using JEOL JEM 1011 (Jeol, Japan) electron microscope (Electron Microscopy Facility, Fondazione Istituto Italiano di Tecnologia, Genoa - Italy) operating with an acceleration voltage of 100 kV and recorded with a 11 Mp fiber optical charge-coupled device (CCD) camera (Gatan Orius SC-1000 - USA). 500.000 BMDMs were seeded into each well of a 6 wells plate. A sterilized borosilicate glass was previously placed into each well; culturing conditions were maintained as described above. Cells were treated overnight with MPN, samples were fixed for 2 hours in 1.5% glutaraldehyde in 0.1 M Sodium Cacodylate buffer (pH 7.4), post fixed in 1% osmium tetroxide in the same buffer and stained overnight with 1% uranyl acetate aqueous solution. Samples were then dehydrated in a graded ethanol series, infiltrated with a series of ethanol/resin solutions and finally embedded in epoxy resin (Epon 812, TAAB). Thin sections were cut with a Leica UC6 ultramicrotome (Leica Microsystems, Germany) equipped with a diamond knife (Diatome).

### Griess assay

100,000 BMDM and 10,000 HUVEC were seeded in 24 well plates (Corning – USA) in 500 µL of media respecting culturing conditions above indicated. Cells were treated varying the MPN concentration from 0 to 0.25 mM equivalent dose of methyl palmitate. After the different time of treatment the media was collected, centrifuged (500 RCF, 5 min, 4 °C) and analyzed.

### Intracellular Ca^2+^ concentration

65.000 BMDMs were seeded into each well of a µ-Slide 8 Well high (ibidi GmbH – Germany) maintaining culturing conditions, as described above. For monitoring intracellular calcium concentration Fluo-4AM, cell permeant (Thermo Fisher – USA) was added to the media according to vendor indication. 1 day after the seeding cells were treated with MPN. For this treatment 1 batch 1× of MPN was dispersed in PBS and sonicated right before the use, 7.5 µL of the suspension was added to the media. Cells were imaged right after by time lapse microscopy using a Nikon Eclipse-Ti-E microscope (Nikon Instrument Inc. – Japan) under controlled environmental conditions. Movies were acquired at a frame rate of 12 fpm using a 40× lens. The events registered from 3 cells were acquired.

### Raster-scanning optoacoustic mesoscopy (RSOM)

Nude mice at 6-8 weeks of age were obtained from Jackson. The tumor-bearing cohort were implanted subcutaneously, between the flank and the dorsal side of the mouse, with 2 million CT26 cells resuspended in 100 µL of PBS. Once tumors reached 500-1000mm3, mice were randomized into PBS or MPN injection groups. For RSOM imaging, mice were placed in an isoflurane chamber for 5 minutes before being positioned on the RSOM imaging bed with continuous isoflurane and oxygen flow. Ultrasound gel was applied to the mouse in the region designated for RSOM imaging. The RSOM imaging head containing the laser, water bed, and transducer were placed on the dorsal side of the mouse containing tumors or healthy skin (no tumor). Once an appropriate area and depth was set with the imaging head, images were then acquired before MPN or PBS injection. Mice were then injected with either 300 µL of PBS or MPN (resuspended in PBS) and image acquisition was initiated at 1-minute post injection (T1). Additional images were acquired every 5 minutes post T1 until the 20-minute timepoint.

The RSOM Viewer program was used to complete motion correction and spectral unmixing on the RSOM images. Spectral unmixing removes noise from melanin and air bubbles and allows for the imaging of oxygenated and deoxygenated blood, as well as high and low frequency wavelengths, corresponding to narrow or constricted vessels or wider/dilated vessels, respectively.

For vessel width analysis, 17 – 20 vessel regionss were chosen at random on blinded images of PBS or MPN injected mice, with the only condition being that there was little background noise surrounding the region of the vessel being analyzed. ImageJ was used to create a stack of the T1 and T4 images, and the plot profiler tool was used to plot the gray value across the length of a 12-pixel line drawn over the vessel/ region of interest. The maximal gray value is then located at the center of the 12-pixel line and is used to normalize all other gray values, and generate a bell curve representing normalized gray value against pixel length, with the largest gray value equaling 1, once normalized. A python script was used to interpolate the pixel values (x1 and x2) at full width half maximum (y = 0.5) for T4 and T1. The change in vessel width (pixel change) from T4 to T1 was then determined by calculating (T4x2- T4x1) – (T1x2 – T1x1), referred to here as ΔΔX . The python script used to calculate the ΔΔX between T1 and T4 timepoints for all the vessel regions analyzed, can be found at: https://github.com/liz-lemon18/Delta_Delta_X/blob/main/delta_delta_x.py. The pixel-to-micron conversion for vessel width measurements was based on specifications provided by iThera for the RSOM P50 Explorer, where 1 pixel corresponds to 20 µm.

Vessels which appeared anew in the T4 time point but were not visible in the T1 time point could not be analyzed, since a plot profile could not be generated for non-visible vessels in T1. Therefore, this analysis focuses mainly on vessels whose change in width could be measured by the plot profile function in T4 vs T1.

### ^89^Zirconium- labeled Feraheme Biodistribution Analysis

6-8 weeks old BALB/c mice obtained from Taconic were implanted in the flank region with 2 million CT26 cells that were suspended in 100 µL of phosphate buffered saline. Once tumors reached a volume between 250 – 1,000 mm^3^, mice were randomized and placed into PBS or MPN pre-injection groups. For injections, mice were placed into a cage for heating for 5 minutes, after which mice were injected with 300uL of PBS or MPN via tail vein injection and then placed in a second cage for a five-minute recovery period. Mice were then anesthetized by using isoflurane for 5 minutes, immediately after which 100µL of ^89^Zr-FH was injected retro-orbitally. 6 hours post injection, mice were imaged using the Siemens Inveon PET/CT imager. Image processing was completed using the Inveon Research Workplace software to quantify the PET signal from sagittal slices of 3D maximum intensity projections (MIPs) of the CT26 tumor mass in PBS and MPN pre-injected mice, using the paintbrush tool to highlight the region of interest (ROI). Mice with tumors of equivalent masses, resulting in near identical average tumor mass and variability among the two pre-treatment groups, were compared for this experiment (n = 4). The mean intensity of the ROIs was then divided by the volume of the tissue analyzed in the 3D MIP to obtain volume-normalized mean intensities for each group.

For gamma count analysis, mice were sacrificed at 24 hours post ^89^Zr-FH injection and organs were collected into pre-weighed 5mL plastic capped tubes. Each sample was read for 60 seconds on the gamma counter to obtain counts per minute (CPM) and an ^89^Zr standard curve was used to convert CPM to activity in microcuries (µCi). After decay correction calculations were completed, data was reported as percent injected dose per gram of tissue (%ID/g) after weighing 5 mL plastic capped tubes once again, post organ harvesting. Percent change and the associated error was calculated using the percent change equation (shown below) in which the averaged PBS 6-hour region-of-interest (ROI) or 24 hour %ID/g value for the CT26 tumor mass was subtracted from each MPN ROI or %ID/g value:

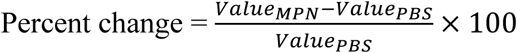

### Intratumoral Dextran Accumulation

. BALB/C mice of 6-8 weeks of age were obtained from Taconic and were subcutaneously implanted in the flank with 2e6 CT26 cells in 100 µL of PBS. Once tumors reached ∼ 400 – 2,000 mm^3^, mice were randomized in MPN or PBS pre-injection groups (5 mice per group). One mouse was set aside for PBS only or MPN only negative control injections. 300 µL of PBS or MPN was injected via tail vein, after which, mice were anesthetized with isoflurane and 100 µL of Dextran-Oregon Green (0.83 mg/mouse) was injected retro-orbitally, within 15-20 minutes of MPN or PBS injection. 24-hour later, mice were sacrificed and tumors were collected and fixed in 4% paraformaldehyde for 1 week at 4 °C. Tumor tissues were then washed twice with PBS without Calcium and Magnesium and incubated in a 40% sucrose and PBS solution for 5 days. Tissues were then embedded in OCT and cut into 10 µm slices and mounted onto microscope slides. Slides were imaged and scanned using a 488 nm laser. Tumor tissue harvested from mice that received the PBS or MPN pre-injection but no Dextran-Oregon Green injection (negative control) were used as negative control and for background subtraction. Slice analysis was conducted by calibrating 2 thresholds on the green channel, 1 permissive for measuring the slide area and one more restrictive to indicate the signal coming from the dextran. The positive area fraction (%) was calculated by dividing the area occupied by the dextran signal and the total area of the slide. For this experiment n=9 animals per group were considered with n>12 slide replicates per animal.

### Blood tests

Healthy immunocompetent C57BL/6 mice were treated with MPN once with a high dose of methyl palmitate = 3.75 mg / 20 g, or twice per week (for 2 weeks) with a lower dose of methyl palmitate = 0.94 mg / 20 g. The injections were performed in the tail vein. 24h after the last treatment, the blood was withdrawn by the retro-orbital plexus by using a heparinized glass capillary (Sigma-Aldrich, USA). For the hematology tests the blood of each animal was transferred into a Microvette CB 300 EDTA K2E (Sarstedt, Germany). For the biochemistry analyzes the blood of each animal was transferred into an eppendorf tube 1.5 mL (Eppendorf, Germany) and centrifuged for 10 min at 10,000 rpm at room temperature. Serum was isolated and stored at -80 °C before proceeding to the analyses. The tests were run at the “Mouse Clinic” of Ospedale San Raffaele in Milan.

### Histology Analyses

Healthy immunocompetent C57BL/6 mice were treated with MPN once with a high dose of methyl palmitate = 3.75 mg / 20 g, or twice per week (for 2 weeks) with a high and a low dose of methyl palmitate: 3.75 mg / 20 g and 0.94 mg / 20 g, respectively. The injections were performed in the tail vein. 24h after the last treatment mice were sacrificed by using CO_2_, the same animal used for blood tests were used for histology (plus the additional group missing in the other analysis). Main organs were collected and washed in PBS. For the liver, the lobe 2 was used for the analyses. Kidneys were split in half following the longer axis. For the lungs, the left lung was used for the analyzes. Hearth was transversally split in two halves having 1 atrium and 1 ventricle per part. Spleen was maintained integer, and no cut was performed. All the samples from one animal were isolated into a single glass vial after an additional PBS wash. Glass vials were kept in ice and PFA 4% was added in order to fully cover all the sample. Vials were kept at 4 °C for 48h, PFA 4% was removed and 3 PBS washes were performed. After the third wash the PBS was completely removed and ethanol 70% was added, allowing all the samples to be immersed. Each organ was then disposed into a single bio cassette for embedding. All the cassettes were collected into 500 mL jars and shipped at room temperature to the “Mouse Clinic” of Ospedale San Raffaele in Milan and processed there.

## References

1. Blanco, E., Shen, H. & Ferrari, M. Principles of nanoparticle design for overcoming biological barriers to drug delivery. Nat. Biotechnol. 33, 941–951 (2015).

2. Palomba, R. et al. Boosting nanomedicine performance by conditioning macrophages with methyl palmitate nanoparticles. Mater. Horiz. 8, 2726–2741 (2021).

3. Ruge, C. A. et al. Uptake of nanoparticles by alveolar macrophages is triggered by surfactant protein A. Nanomedicine 7, 690–693 (2011).

4. Baumann, D. et al. Complex Encounters: Nanoparticles in Whole Blood and Their Uptake Into Different Types of White Blood Cells. Nanomedicine 8, 699–713 (2013).

5. Jain, R. K. Transport Phenomena in Tumors. in Advances in Chemical Engineering (ed. Wei, J.) vol. 19 129–200 (Academic Press, 1994).

6. Chauhan, V. P., Stylianopoulos, T., Boucher, Y. & Jain, R. K. Delivery of Molecular and Nanoscale Medicine to Tumors: Transport Barriers and Strategies. Annu. Rev. Chem. Biomol. Eng. 2, 281–298 (2011).

7. Stylianopoulos, T. et al. Coevolution of Solid Stress and Interstitial Fluid Pressure in Tumors During Progression: Implications for Vascular Collapse. Cancer Res. 73, 3833–3841 (2013).

8. Siemann, D. W. The unique characteristics of tumor vasculature and preclinical evidence for its selective disruption by Tumor-Vascular Disrupting Agents. Cancer Treat. Rev. 37, 63–74 (2011).

9. Martin, J. D., Seano, G. & Jain, R. K. Normalizing Function of Tumor Vessels: Progress, Opportunities, and Challenges. Annu. Rev. Physiol. 81, 505–534 (2019).

10. Choi, Y. & Jung, K. Normalization of the tumor microenvironment by harnessing vascular and immune modulation to achieve enhanced cancer therapy. Exp. Mol. Med. 55, 2308–2319 (2023).

11. Chauhan, V. P. et al. Normalization of tumour blood vessels improves the delivery of nanomedicines in a size-dependent manner. Nat. Nanotechnol. 7, 383–388 (2012).

12. Dasgupta, A. et al. Nonspherical ultrasound microbubbles. Proceedings of the National Academy of Sciences 120, e2218847120 (2023).

13. Xu, Q. et al. NO-dependent vasodilation and deep tumor penetration for cascade-amplified antitumor performance. Journal of Controlled Release 347, 389–399 (2022).

14. Huang, Z., Fu, J. & Zhang, Y. Nitric Oxide Donor-Based Cancer Therapy: Advances and Prospects. J. Med. Chem. 60, 7617–7635 (2017).

15. Lee, Y.-C. et al. Methyl Palmitate: A Potent Vasodilator Released in the Retina. Invest. Ophthalmol. Vis. Sci. 51, 4746–4753 (2010).

16. Wang, N., Kuczmanski, A., Dubrovska, G. & Gollasch, M. Palmitic Acid Methyl Ester and Its Relation to Control of Tone of Human Visceral Arteries and Rat Aortas by Perivascular Adipose Tissue. Front. Physiol. Volume 9-2018, (2018).

17. Hamed, A. B., Mantawy, E. M., El-Bakly, W. M., Abdel-Mottaleb, Y. & Azab, S. S. Putative anti-inflammatory, antioxidant, and anti-apoptotic roles of the natural tissue guardian methyl palmitate against isoproterenol-induced myocardial injury in rats. *Futur*. J. Pharm. Sci. 6, 31 (2020).

18. Palomba, R. et al. Modulating Phagocytic Cell Sequestration by Tailoring Nanoconstruct Softness. ACS Nano 12, 1433–1444 (2018).

19. Decuzzi, P., Pasqualini, R., Arap, W. & Ferrari, M. Intravascular Delivery of Particulate Systems: Does Geometry Really Matter? Pharm. Res. 26, 235–243 (2009).

20. Champion, J. A. & Mitragotri, S. Role of target geometry in phagocytosis. Proceedings of the National Academy of Sciences 103, 4930–4934 (2006).

21. Hayashi, Y. et al. Differential Nanoparticle Sequestration by Macrophages and Scavenger Endothelial Cells Visualized in Vivo in Real-Time and at Ultrastructural Resolution. ACS Nano 14, 1665–1681 (2020).

22. Gustafson, H. H., Holt-Casper, D., Grainger, D. W. & Ghandehari, H. Nanoparticle uptake: The phagocyte problem. Nano Today 10, 487–510 (2015).

23. Huang, Z. et al. Stimulation of Unprimed Macrophages with Immune Complexes Triggers a Low Output of Nitric Oxide by Calcium-dependent Neuronal Nitric-oxide Synthase*. J. Biol. Chem. 287, 4492–4502 (2011).

24. Connelly, L., Jacobs, A. T., Palacios-Callender, M., Moncada, S. & Hobbs, A. J. Macrophage Endothelial Nitric-oxide Synthase Autoregulates Cellular Activation and Pro-inflammatory Protein Expression*. Journal of Biological Chemistry 278, 26480–26487 (2003).

25. Haedicke, K. et al. High-resolution optoacoustic imaging of tissue responses to vascular-targeted therapies. *Nat*. Biomed. Eng. 4, (2020).

26. Skubal, M. et al. Vascularized tumor on a microfluidic chip to study mechanisms promoting tumor neovascularization and vascular targeted therapies. Theranostics 15, 766–783 (2025).

27. Trujillo-Alonso, V. et al. FDA-approved ferumoxytol displays anti-leukaemia efficacy against cells with low ferroportin levels. Nat. Nanotechnol. 14, 616–622 (2019).

28. Stater, E. P. et al. Translatable Drug-Loaded Iron Oxide Nanophore Sensitizes Murine Melanoma Tumors to Monoclonal Antibody Immunotherapy. ACS Nano 17, 6178–6192 (2023).

29. Zanganeh, S. et al. Iron oxide nanoparticles inhibit tumour growth by inducing pro-inflammatory macrophage polarization in tumour tissues. Nat. Nanotechnol. 11, 986–994 (2016).

30. Kaittanis, C. et al. Environment-responsive nanophores for therapy and treatment monitoring via molecular MRI quenching. Nat. Commun. 5, 3384 (2014).

31. Yuan, H. et al. Heat-induced radiolabeling and fluorescence labeling of Feraheme nanoparticles for PET/SPECT imaging and flow cytometry. Nat. Protoc. 13, 392–412 (2018).

32. Boros, E., Bowen, A. M., Josephson, L., Vasdev, N. & Holland, J. P. Chelate-free metal ion binding and heat-induced radiolabeling of iron oxide nanoparticles. Chem. Sci. 6, 225–236 (2015).

33. Battelli, M. G., Bortolotti, M., Polito, L. & Bolognesi, A. The role of xanthine oxidoreductase and uric acid in metabolic syndrome. Biochimica et Biophysica Acta (BBA) - Molecular Basis of Disease 1864, 2557–2565 (2018).

34. Fukahori, M., Ichimori, K., Ishida, H., Nakagawa, H. & Okino, H. Nitric Oxide Reversibly Suppresses Xanthine Oxidase Activity. Free Radic. Res. 21, 203–212 (1994).

35. J, M. A. & A, B. K. Uric acid is closely linked to vascular nitric oxide activity. JACC 38, 1850–1858 (2001).

