## Supplemental Data for "Enhancing Tumor Perfusion and Nanomedicine Delivery via Endogenous Nitric Oxide Release by Methyl Palmitate Nanoparticles"

#### Supplemental Information


- Roberto Palomba[1 ♦], Elizabeth Isaac[2 ♦], Raffaele Spanò[1], Federica Piccardi[1], Benedict McLarney[2], Elana Apfelbaum[2], Nermin Mostafa[2], Charlene Hsu[2], Jan Grimm [2, 3 #] and Paolo Decuzzi [1,4 #]

- 1 Laboratory of Nanotechnology for Precision Medicine – Fondazione Istituto Italiano di Tecnologia, Via Morego 30, 16163, Genova, Italy

- 2 Molecular Pharmacology Program, Memorial Sloan Kettering Cancer Center, 1275 York Avenue, New York, NY, USA

- 3 Department of Radiology, Memorial Sloan Kettering Cancer Center, 1275 York Avenue, New York, NY, USA

- 3 School of Medicine/Division of Oncology, Center for Clinical Sciences Research, Stanford University, 269 Campus Drive, Stanford, CA 94305 - USA

- ♦ Shared first authorship

- # corresponding authors

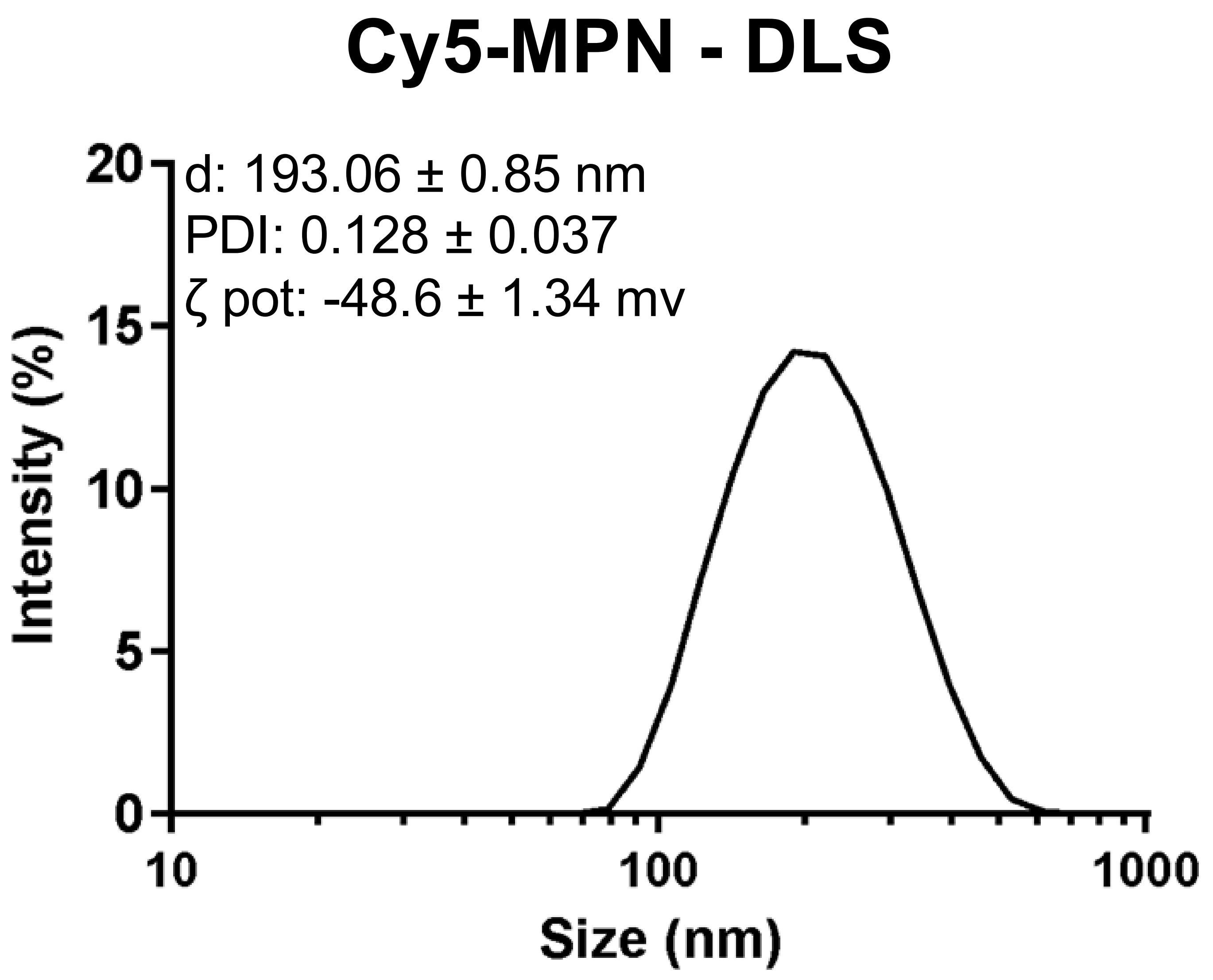


Supplementary Information 1: Dynamic light scattering (DLS) analysis showing the hydrodynamic size distribution of Cy5-MPN, including diameter (d), polydispersity index (PDI), and surface electrostatic potential (ζ).

### Raster Scanning Optoacoustic Mesoscopy of Healthy Tissue

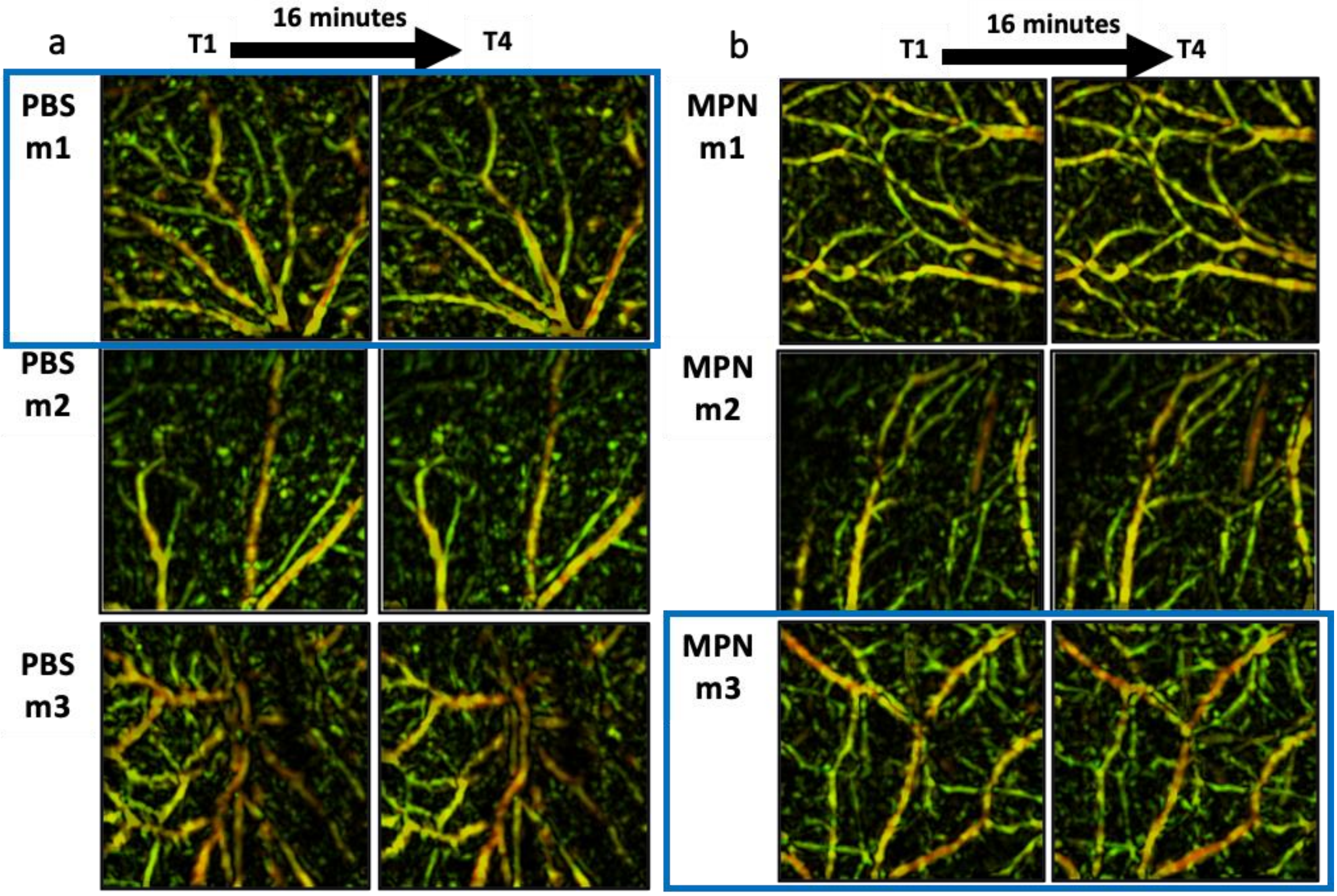


Supplementary Information 2: Raster Scanning Optoacoustic Mesoscopy (RSOM) of mouse healthy (non-tumor) tissue after injection with PBS or MPN at timepoints T1 (1 minute post injection) or T16 ( 16 minutes post injection) in n=3 mice. Representative images chosen for use in the main text are outlined with blue boxes.

**Raster Scanning Optoacoustic Mesoscopy of Tumor Tissue**

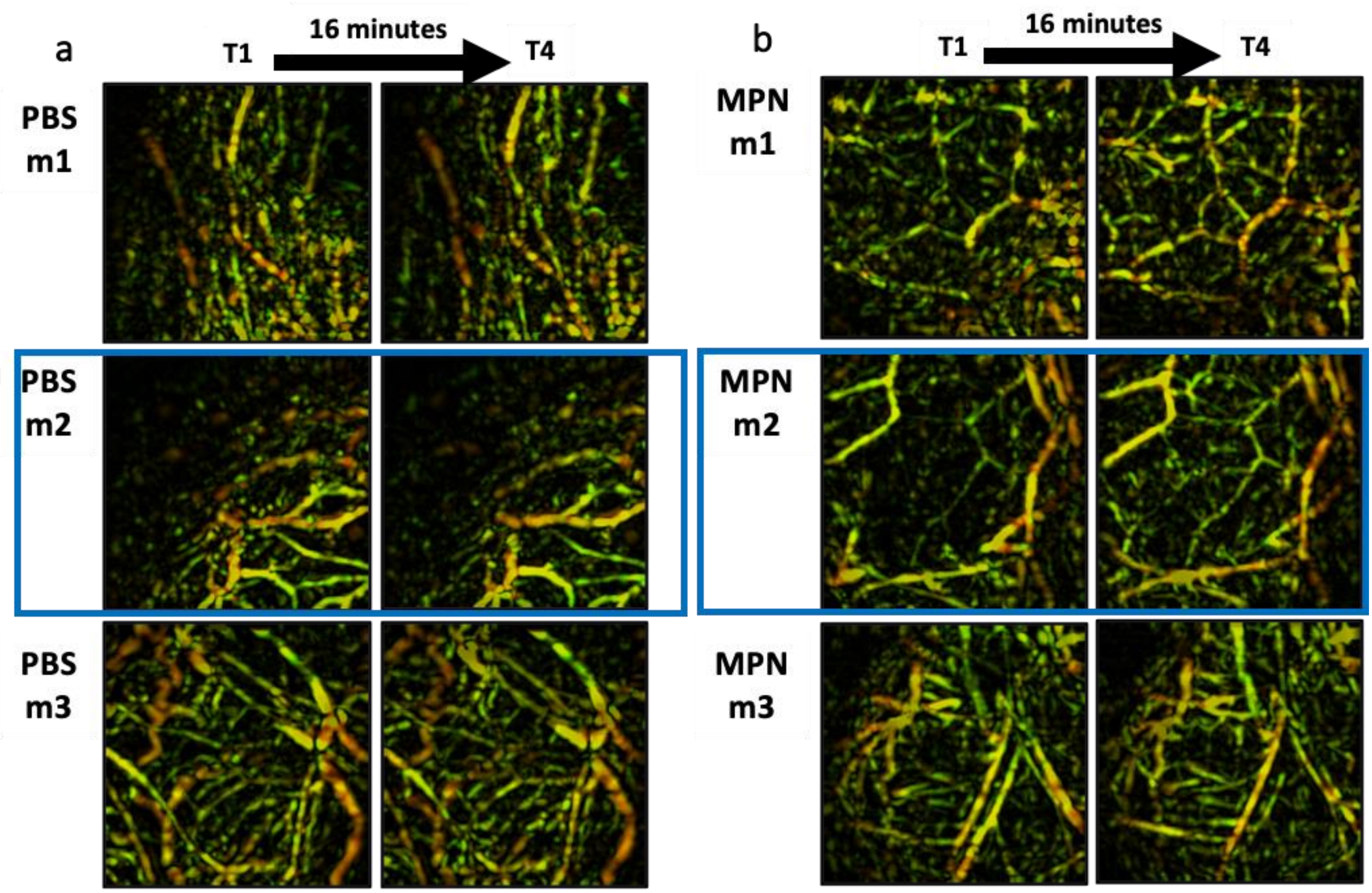


Supplementary Information3: Raster Scanning Optoacoustic Mesoscopy (RSOM) of mouse CT26 tumor tissue after injection with PBS or MPN at timepoints T1 (1 minute post injection) or T16 ( 16 minutes post injection) in n=3 mice. Representative images chosen for use in the main text are outlined with blue boxes.

#### PET/CT images of 89Zr-FH injected mice pre-injected with PBS or MPN

a

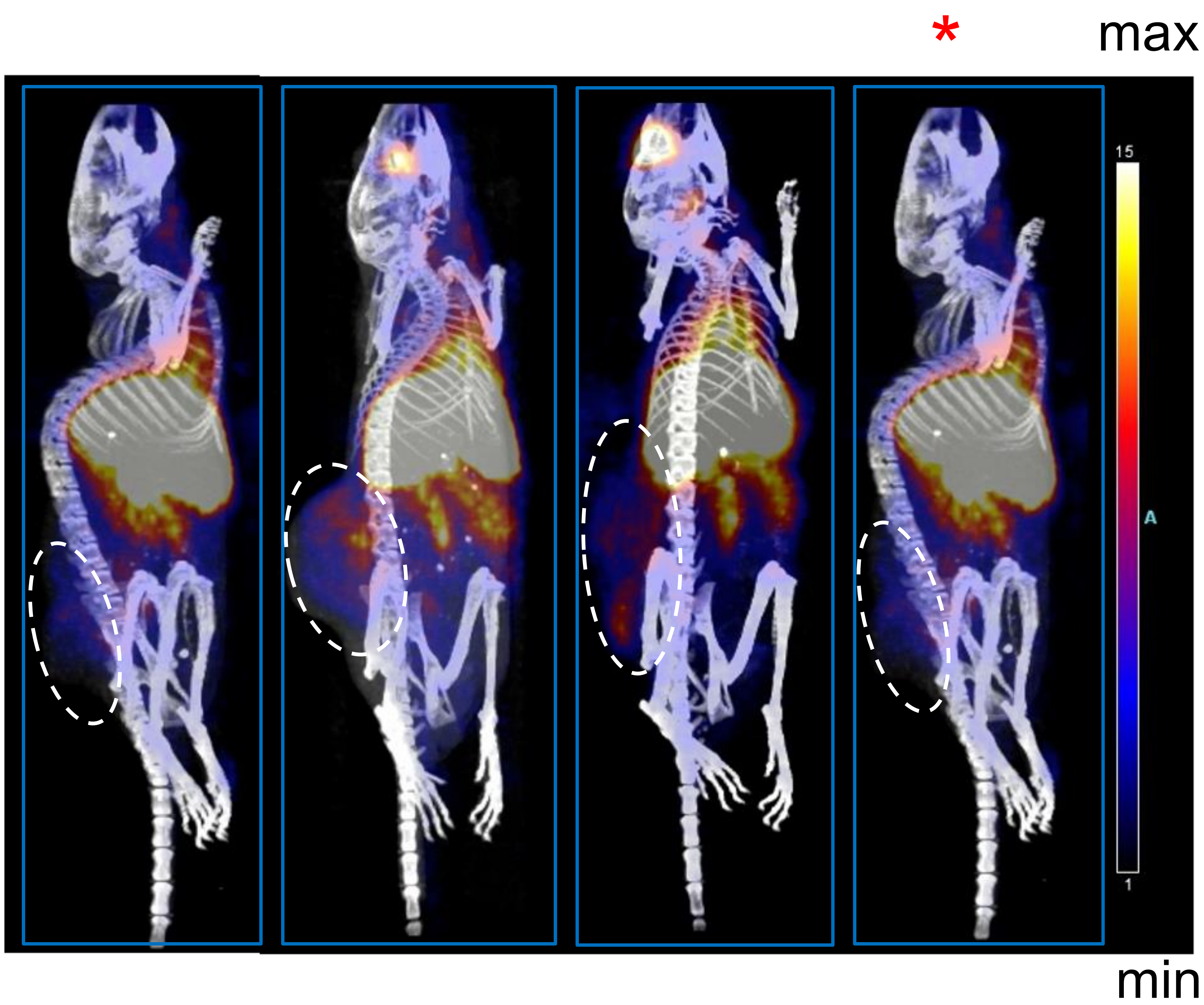


b

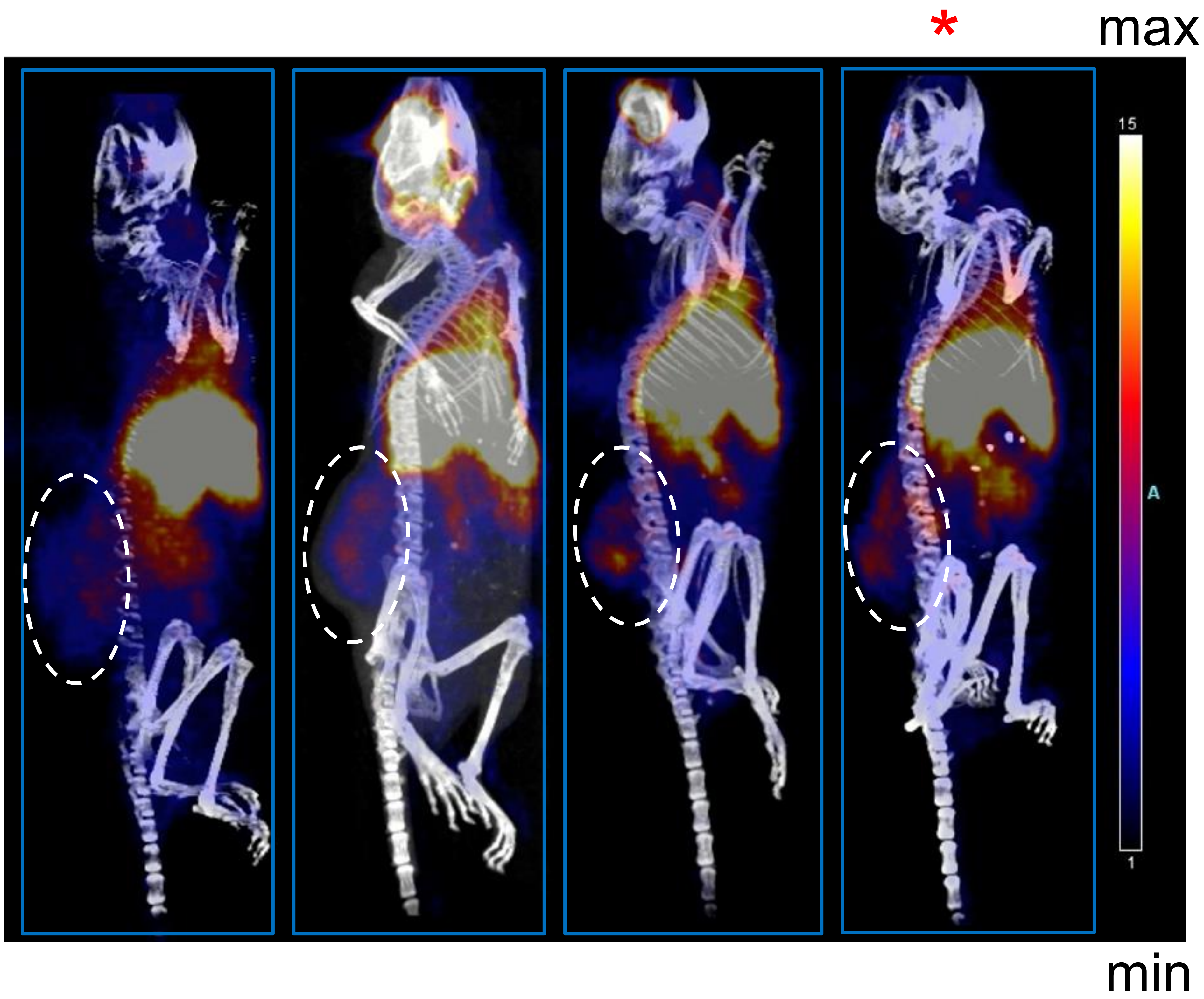


Supplementary Information 4: PET/CT images of n=4 mice acquired 6-hours post injection with 89Zr-FH. Mice chosen as representative images are marked with an asterisk (*) symbol. Tumors are indicated with dashed circles.

### Serological Blood Parameters

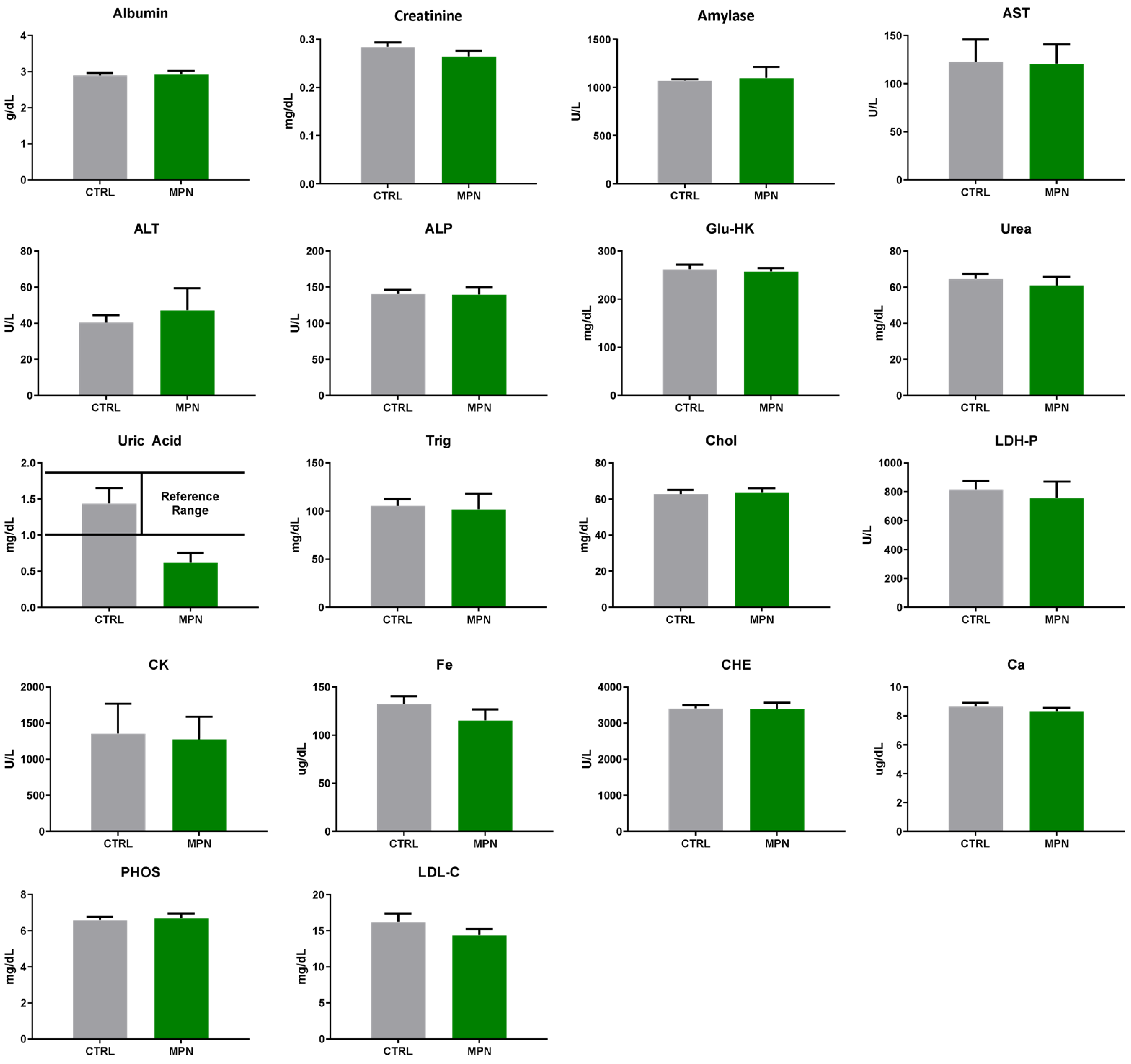


Supplementary Information 5: serological blood parameters in C57BL/6 mice that received a single high dose of MPN corresponding to 3.75 mg of methyl palmitate per 20 g body weight, administered via tail-vein injection.

### Hematological Blood Parameters

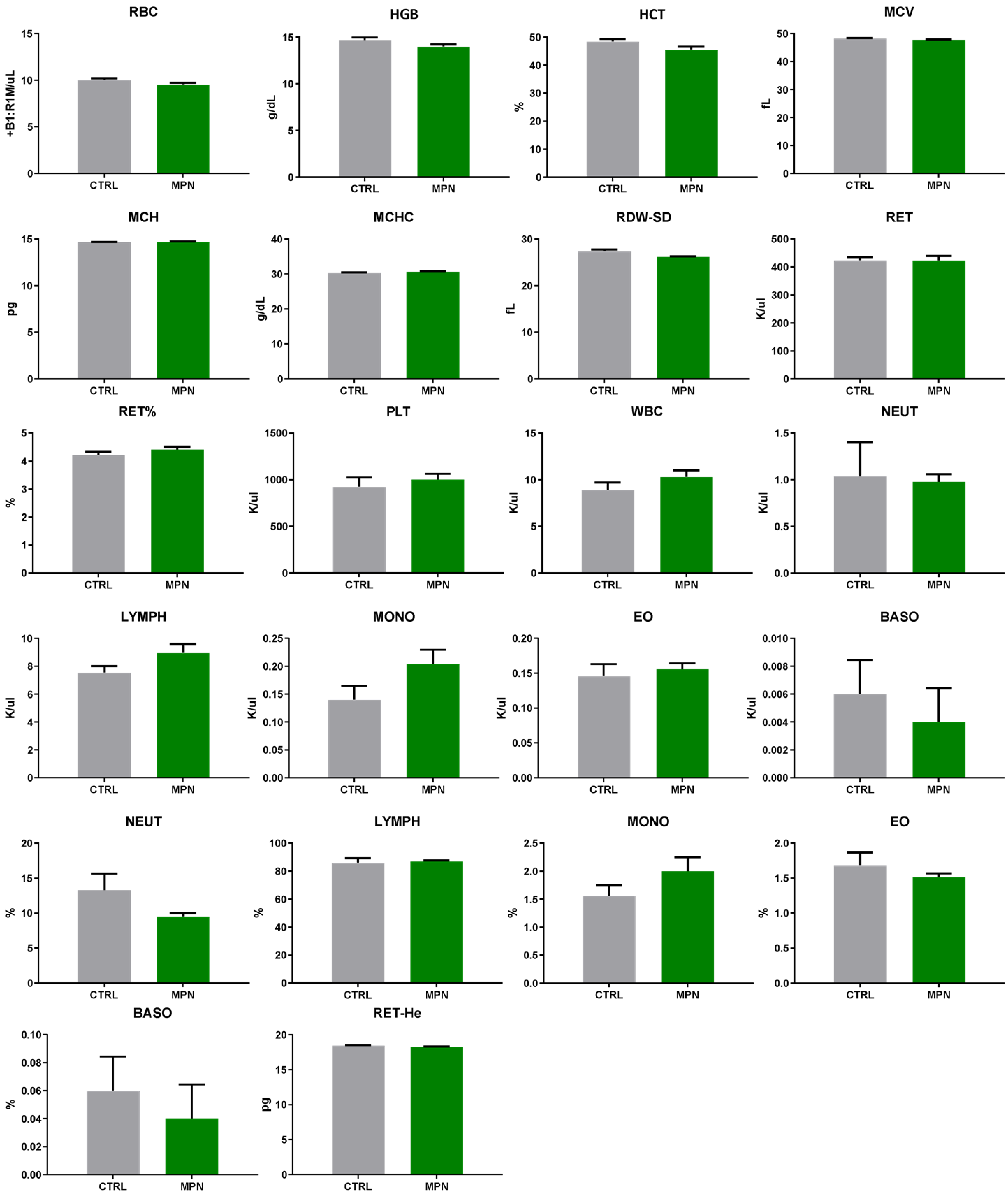


Supplementary Information 6: hematological parameters in C57BL/6 mice that received a single high dose of MPN corresponding to 3.75 mg of methyl palmitate per 20 g body weight, administered via tail-vein injection.

### Serological Blood Parameters

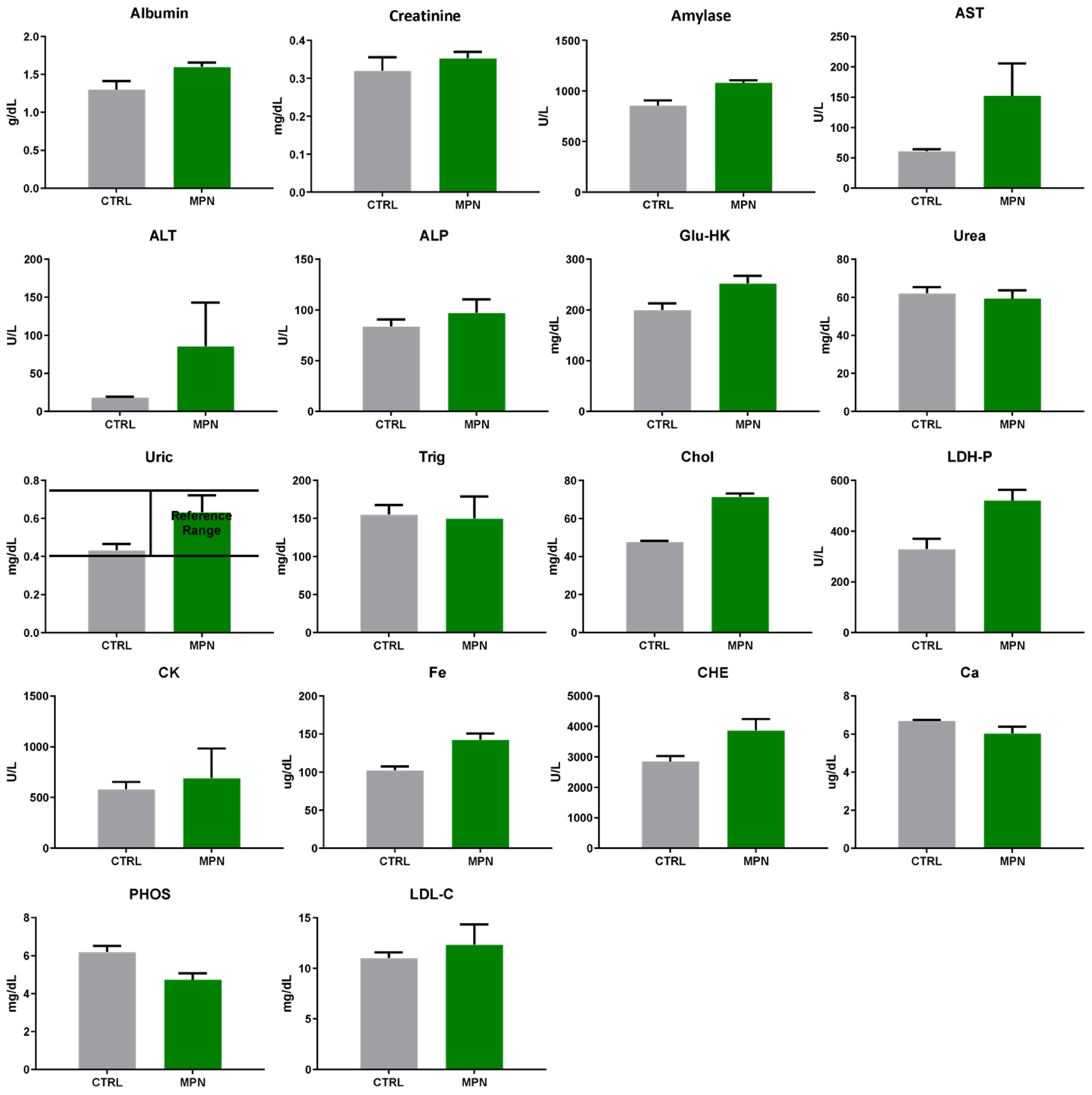


Supplementary Information 7: serological blood parameters in C57BL/6 mice received four injections of MPN administered twice per week over a 2-week period at a lower dose equivalent to 0.94 mg of methyl palmitate per 20 g body weight, administered via tail-vein injection.

### Hematological Blood Parameters

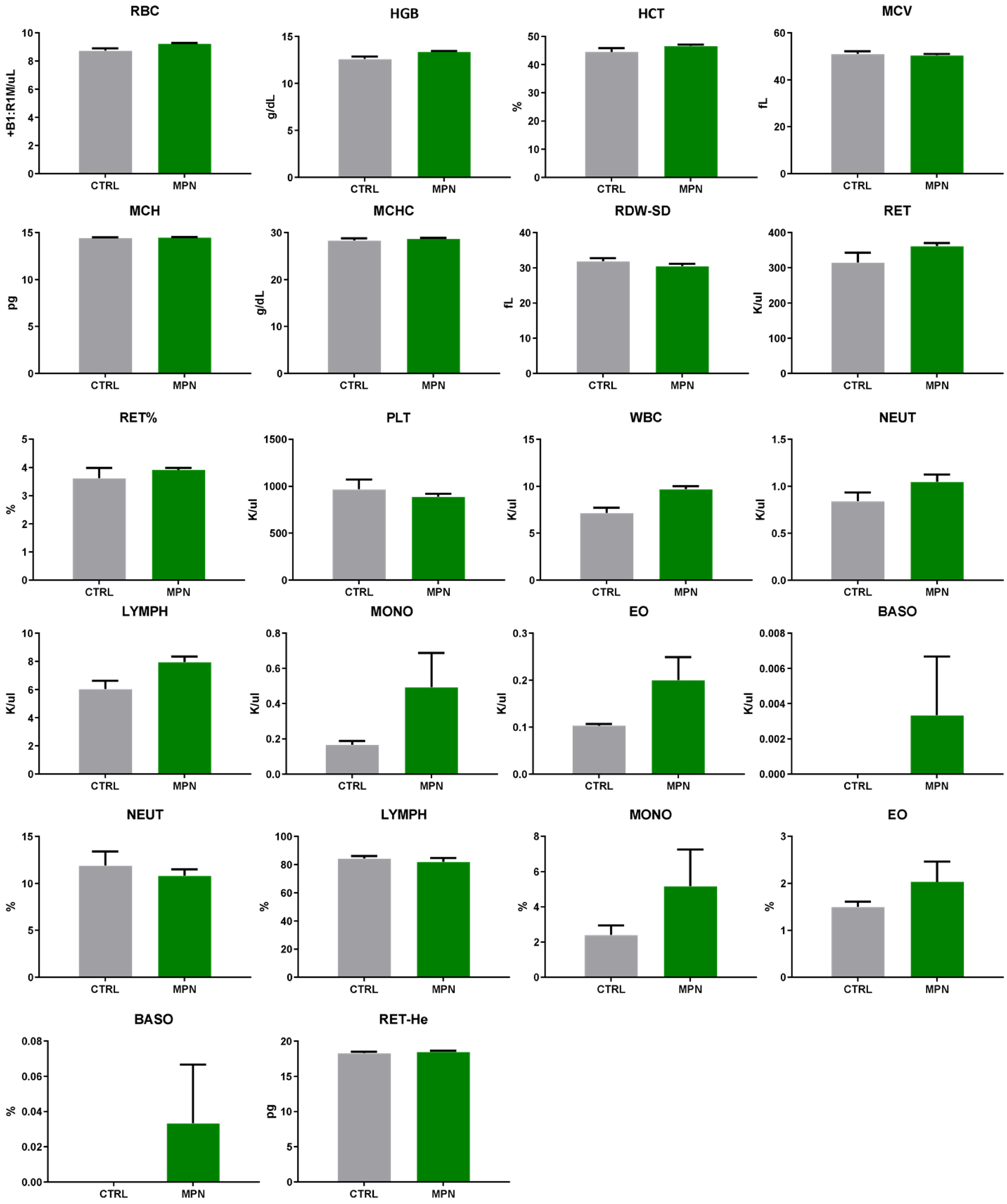


Supplementary Information 8: hematological parameters in C57BL/6 mice received four injections of MPN administered twice per week over a 2-week period at a lower dose equivalent to 0.94 mg of methyl palmitate per 20 g body weight, administered via tail-vein injection.
