## Supplementary figures and images for "Enhancing Tumor Perfusion and Nanomedicine Delivery via Endogenous Nitric Oxide Release by Methyl Palmitate Nanoparticles"

### Tables 1 & 2

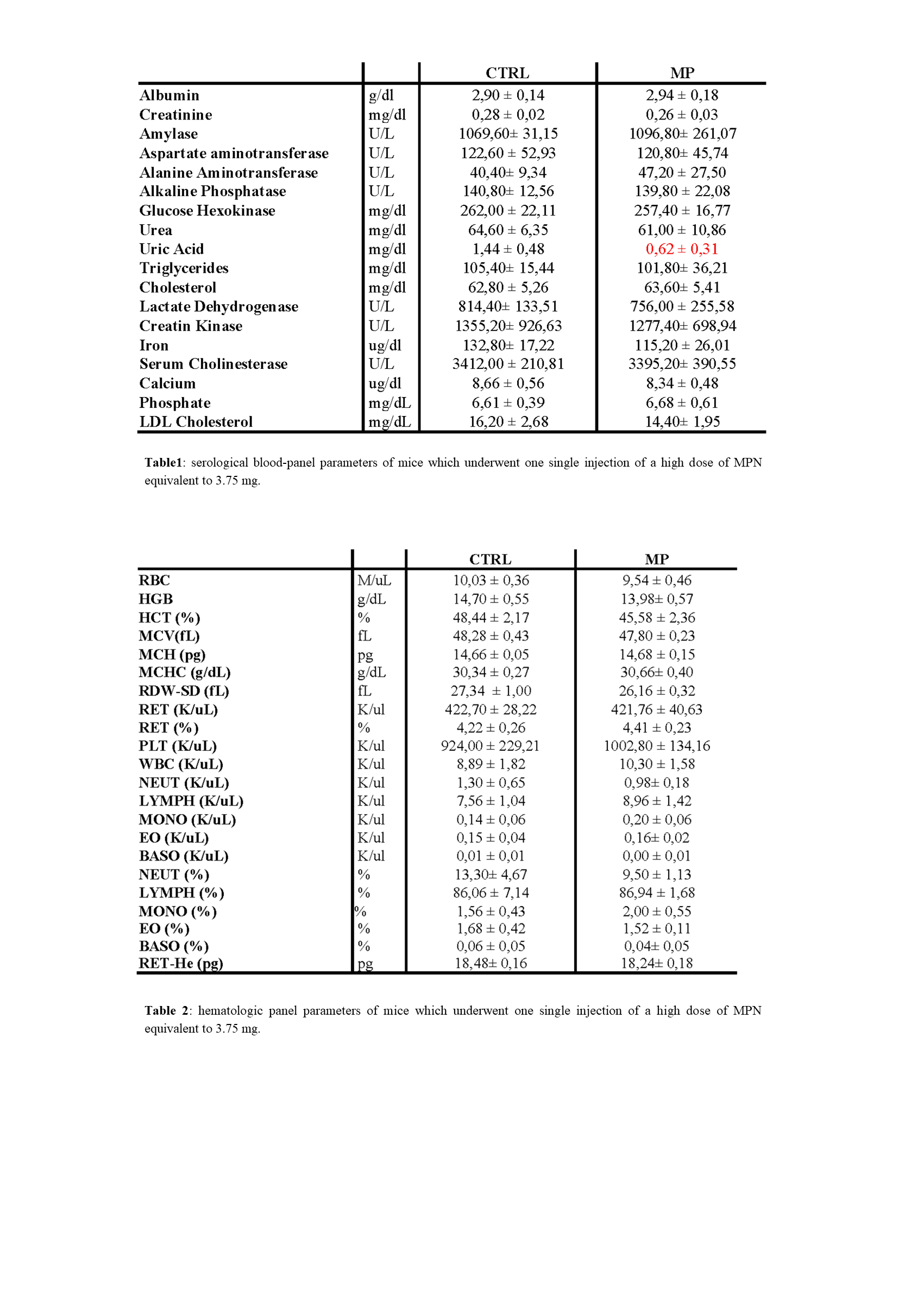

### Tables 3 &4

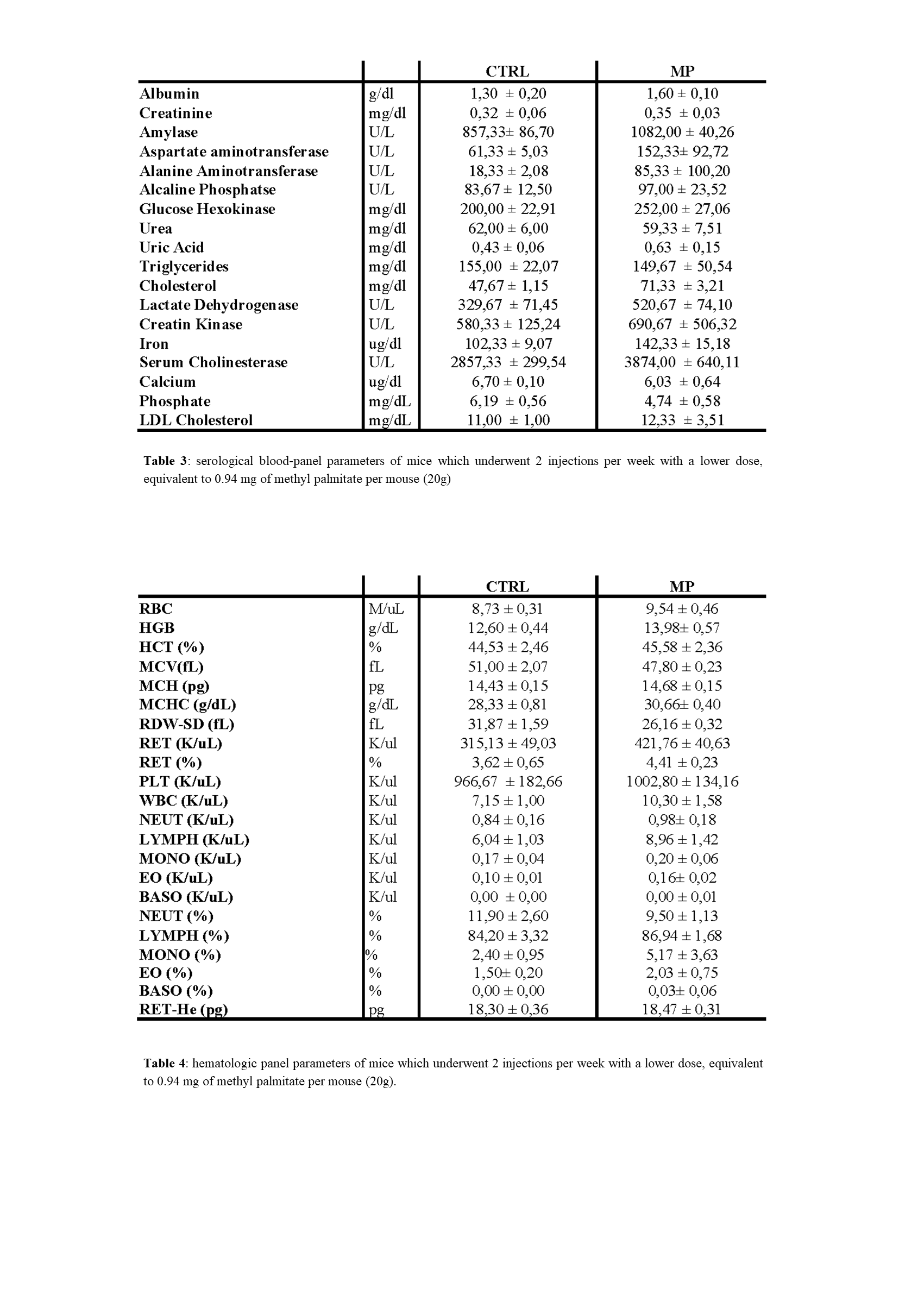
